# Insect-derived long non-coding RNAs function as epigenetic effectors to reprogram plant immunity

**DOI:** 10.1101/2025.09.21.677575

**Authors:** Dong Wen, Shan Jiang, Zhuangzhuang Qiao, Chi Liu, Jun Wu, Rong Hu, Yeqi Zhu, Yueping He, Weihua Ma, Hongxia Hua, Yazhou Chen

## Abstract

Cross-kingdom RNAs are emerging as critical mediators of interspecies interactions, yet the functions of long RNAs such as mRNAs and long non-coding RNAs (lncRNAs) in recipient organisms remain largely unexplored. Here, we show that the brown planthopper (*Nilaparvata lugens*, BPH), a major rice pest, translocates mRNAs and lncRNAs into rice plants, where they migrate systemically from feeding sites to distal tissues. Compared with BPH mRNAs, *BPH Salivary gland Cross-kingdom LncRNA* (*BSCL*s) exhibit markedly higher stability in rice. Among them, *BSCL1* functions as a virulence factor that promotes BPH feeding and reproduction by suppressing host defense. Mechanistically, *BSCL1* associates with the HIRA histone chaperone complex and displaces histone H3.3 from the promoters of transcription factors, including bHLH genes central to jasmonic acid signaling, thereby repressing transcriptional immunity. Our results identify BSCLs as systemic, RNA-based effectors that reprogram host defense at the epigenetic level, revealing a previously unrecognized mode of insect-mediated manipulation of plant immunity and highlighting lncRNAs as cross-kingdom regulators.

**Significance Statement:** Plant–herbivore interactions are traditionally viewed as battles over nutrients and defense signaling, mediated largely by proteins and small RNAs. Here, we demonstrate that long non-coding RNAs from brown planthopper saliva are translocated into rice, migrate systemically, and function as epigenetic effectors that suppress key transcription factors in jasmonic acid–mediated defense. Unlike insect mRNAs, these lncRNAs persist in the plant and reprogram immunity by interfering with histone deposition. This discovery uncovers a new class of mobile, RNA-based virulence factors, expands our understanding of cross-kingdom regulation, and suggests innovative strategies for pest control targeting RNA effectors.

## Introduction

Cross-kingdom RNAs are RNA molecules of an organism that traffic and function in recipient organisms that belong to different biological kingdoms. Among these, cross-kingdom RNA interference (RNAi) phenomena that involve small RNAs (sRNAs) have been widely observed across various parasite-host interactions (1–5). Those sRNAs traffic across the organisms and influence gene expression in the recipients, which reveals the critical roles of cross-kingdom RNAs in these interactions.

In contrast, far less attention has been given to the functions of long RNA species such as mRNAs and long non-coding RNAs (lncRNAs) in recipient organisms, although they have been found to be intensively trafficked during parasite-host interactions. For example, plant phloem mRNAs have been found to traffic bidirectionally between parasite dodders and various host plants (6, 7). Arabidopsis mRNAs are delivered into the fungus and then translated by the fungal cells, leading to the reduction of fungal infection (8). *Cryptosporidium parvum* lncRNAs were selectively delivered into the nuclei of human intestinal epithelial cells during infection (9). Aphid lncRNAs are translocated into divergent host plants and systematically migrate in the plants and promote aphid colonization (10). These studies suggest that long RNAs play essential roles in mediating parasite-host interactions; however, their function in the hosts remains largely unexplored.

Sap-feeding insects such as aphids and planthoppers are excellent models for studying cross-kingdom RNAs. These insects suck the contents from the xylem and/or phloem tissues, the main components of plant vascular tissues (11–14). Watery saliva secreted by insect salivary glands is injected into plant vascular tissues, where salivary molecules, including various RNAs, enter and traffic in the plants (5, 10, 13, 15, 16). Moreover, these insects can be easily confined to specific areas of the plant, leaving the rest of the plant intact and uncontaminated, enabling tracking the migration of insect RNAs in the plant by various approaches such as qRT-PCR (5, 10, 16). Additionally, insect-derived RNAs, as foreign RNA species, exhibit low sequence similarity to plant RNAs, making them highly distinguishable and ideal for detection using next-generation sequencing approaches (5, 10).

Here, we used the brown planthopper (*Nilaparvata lugens* Stål, BPH), a major rice pest worldwide (17), to investigate the translocation of insect-derived RNAs into host plants. Our study identified both lncRNAs and mRNAs delivered into rice, with particular focus on salivary gland–derived lncRNAs, termed *BSCL*s (*BPH Salivary gland Cross-kingdom LncRNA*s). These *BSCL*s were translocated into rice and migrated systemically from the feeding site to distal tissues, including leaves and roots. Compared with BPH mRNAs, *BSCLs* exhibited markedly higher stability within rice plants. Among them, *BSCL1* promoted BPH feeding and reproduction by suppressing rice defense responses. *BSCL1* displaced histone complexes from the promoters of transcription factor genes, leading to transcriptional repression of transcription factors such as bHLH, central to jasmonic acid (JA) signaling, thereby weakening host resistance to BPH.

## Results

### Identification and host-responsive expression of salivary lncRNAs in BPH

To annotate lncRNAs in the BPH genome, we assembled strand-specific RNA-seq data from 11 libraries (Table S1), identifying 49,435 transcripts from 47,350 genes (Fig. S1*A*). After filtering, 24,433 putative lncRNAs (22,374 genes) were retained, including 10,268 previously reported and 12,106 newly identified (Fig. S1*A*). Comparative analysis across 28 arthropod genomes showed that most lncRNA genes (57.9%, 12,957) are BPH-specific (Fig. S1 *B* and *C*), suggesting a recent evolutionary origin.

To explore potential functions in BPH–rice interactions, we analyzed lncRNA expression in insects transferred from susceptible TN1 rice to resistant RHT plants. RNA-seq revealed 1,626 down-regulated and 1,291 up-regulated genes (Fig. S1*D*, Dataset S1), including 932 lncRNAs enriched in salivary gland expression (Fig. S1*E*). Notably, 606 differentially expressed lncRNAs were expressed in salivary glands, and most (78.7%, 477) were repressed by resistant plants (Fig. S1 *F* and *G*), implicating salivary lncRNAs in host colonization.

### BPH salivary gland lncRNAs are translocated into rice plants

Previously, we reported that aphid salivary gland lncRNAs are translocated into plants, and lncRNA *Ya1* acted as a virulence factor to promote aphid colonization (10). Thus, we investigated whether BPH salivary gland lncRNAs are secreted into rice via feeding. We conducted cage experiments in which 30 male BPHs—chosen to prevent egg deposition on the plants—were confined to the leaf sheath (designated as feeding sites, FS) of 45-day-old rice plants for 2 days (Fig. 1*A*). Controls were counterparts of untreated plants which the sheath was in an empty cage. Caged tissues were carefully washed to remove visible BPH tissues and were subsequently subjected to RNA-seq analysis. Bioinformatically, adaptor-removed RNA-seq reads were aligned to a merged genome file that the concatenated reference genomes of rice TN1 (18) and BPH (19) (Fig. S2*A*). Expectedly, the vast majority of reads in the replicates of controls and FS were mapped to the merged genome (Dataset S2). To gain more confidence, only uniquely aligned reads were used in the following analysis.

**Fig. 1.**
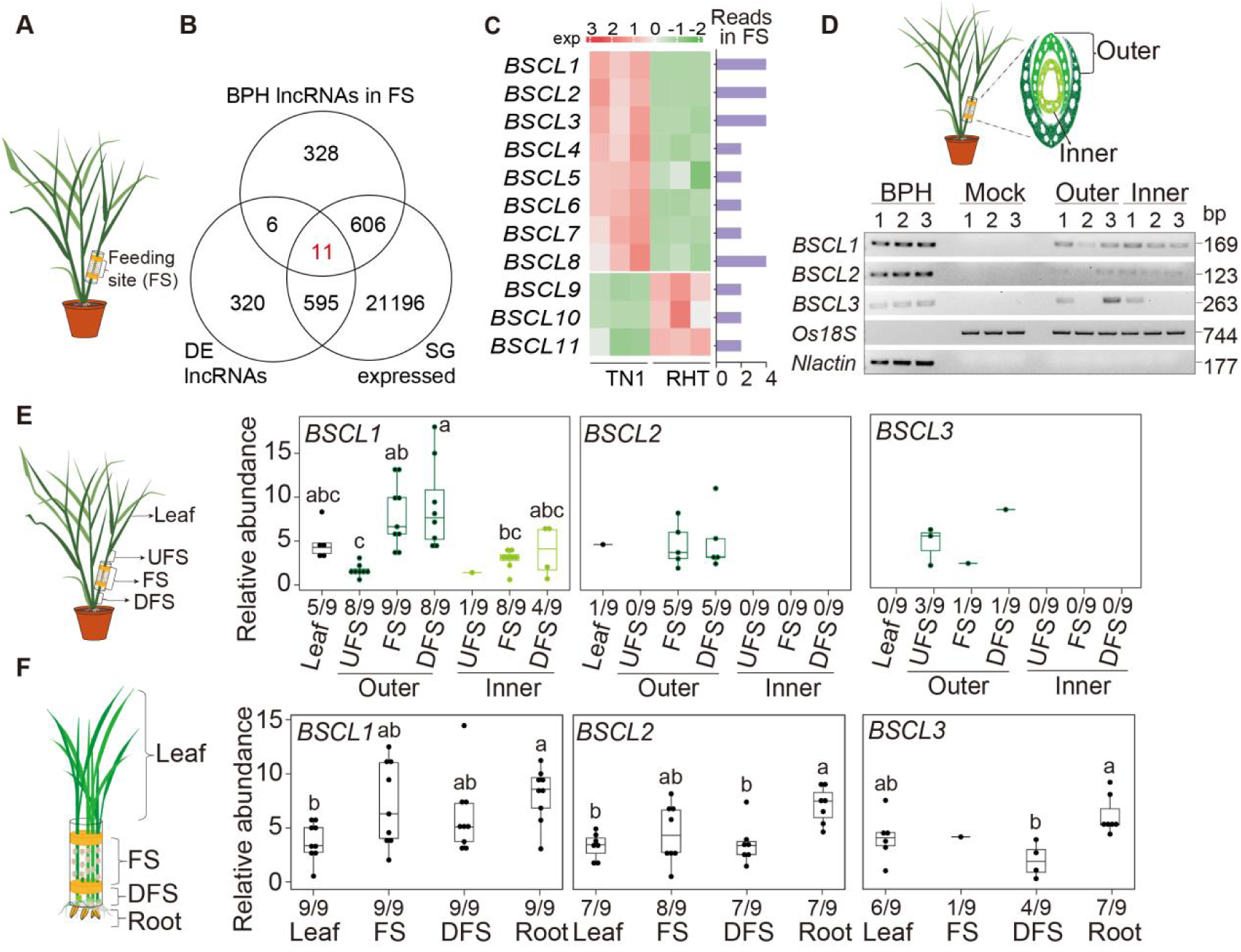
BPH-delivered lncRNAs systematically migrate in rice. (A) Schematic overview of the feeding experiment. FS, the leaf sheath segment directly exposed to BPH feeding. (B) Venn diagram showing the overlap of lncRNAs identified in FS, DE in response to RHT, and expressed in BPH salivary glands. (C) Heatmap of *BSCLs* transcript abundance (TPM values). The bar plot shows the sequencing reads of *BSCLs* detected in FS. (D) Migration of *BSCL1*, *BSCL2*, and *BSCL3* from the outer to inner layers of the FS. A schematic of inner and outer layers is shown on the left. (E) Migration of *BSCL1*, *BSCL2*, and *BSCL3* from FS to the UFS, DFS regions of the sheath, and the leaf blade. A schematic of FS, UFS, and DFS is shown on the left. Box plots indicate the relative abundance of *BSCL1*, *BSCL2*, and *BSCL3* at different sampling sites, determined by qRT-PCR. (F) Migration of *BSCL1*, *BSCL2*, and *BSCL3* to the rice roots. A schematic of leaf blade, FS, DFS and root is shown on the left. Box plots indicate the relative abundance of *BSCL1*, *BSCL2*, and *BSCL3* at different sampling sites, determined by qRT-PCR. Numbers under the boxpot show the detection ratio of *BSCL1, BSCL2* and *BSCL3* at each site. Different lowercase letters indicate statistically significant differences at *P* < 0.05 according to one-way ANOVA followed by Tukey’s multiple comparisons test.

The number of BPH reads in the FS varied largely among the replicates, resulting in different numbers of BPH lncRNAs identified (Dataset S2, Fig. S2*B*). For instance, 3065 uniquely mapped reads from 951 transcripts were identified from one replicate, while fewer than 40 reads were identified in the other two. Thereby, we conducted two more independent experiments with two replicates each. Again, the number of genes detected varied among replicates (20 to 236) because of different numbers of uniquely mapped reads between replicates. Such variation has been observed in other BPH experiments (5). We chose the sample BPH-1-FS from replicate 1 for further analysis since the highest number of BPH RNA reads were detected (244 transcripts with reads ≥ 4, Dataset S3).

Among these 244 transcripts that are likely to be cross-kingdom RNAs (Dataset S3), 11 were expressed in the salivary glands and were also DE between BPH that fed on TN1 and RHT (Fig. 1*B*, Dataset S1), which included 8 down-regulated and 3 up-regulated lncRNA genes (Fig. 1*C*). We found that 10 out of 11 BPH lncRNAs were detected in the FS of rice plants in the repeated cage experiments but were absent from the same RNA samples that weren’t treated with reverse transcriptase and from the counterparts on the control plants (Fig. S2*C*). Therefore, these lncRNAs were named *BPH Salivary gland Cross-kingdom LncRNA* (*BSCL*). Noticeably, the PCR bands of 7 *BSCL*s (*BSCL4-9* and *BSCL11*) were faint unless the PCR products were used as templates for another round of PCR amplification (Fig. S2*C*). Three BSCLs (*BSCL1*, *BSCL2*, *BSCL3*) were detected in more than two tested samples and absent from the controls (Fig. S2*C*).

In conclusion, BPH lncRNAs are translocated into rice plants via feeding, although there is some variation among different individuals.

### BPH *BSCL*s migrate systemically within rice plants

Detection of BPH lncRNAs in the FS of rice sheath prompted us to assess whether these RNAs migrate inside rice plants. BPH feeding occurs mainly at the outer layer of rice stem sheaths, thus the FS tissues were dissected into the outer and inner layers (Fig. 1*D*). The presence of *BSCL1*, *BSCL2*, and *BSCL3* on each layer was examined (Fig. 1*D*). PCR amplification with specific primers detected three *BSCLs* clearly in both layers, suggesting migration of these *BSCL*s in the rice plants.

In addition to the FS, the tissues that were about 2 cm above and below the caged sheath, respectively termed up- and down-near feeding sites (UFS and DFS, Fig. 1*E*), were also harvested. All three *BSCLs* were detected in the outer layers of FS and DFS (Fig. 1*E*). Except for *BSCL2*, both *BSCL1* and *BSCL3* were also found in the outer layers of UFS. Sequences of PCR products of these three lncRNAs obtained from rice sheath further confirmed the translocation of BPH lncRNAs into the rice plants (Fig. S3 *A*-*C*). *BSCL1* was also found in the inner layers of UPS, FS, and DFS (Fig. 1*E*), probably because of its relatively higher expression level.

*BSCL1* abundance in the outer layers ranked from the DFS, FS, then to UFS, which was similar as that in the inner layers (Fig. 1*E*). It indicated that migration of *BSCL*s in the rice plants is likely from top to bottom in most outer sheaths where BPH actually feed on, and from bottom to top in the inner sheaths. The opposite may be achieved through the internodes, where nutrients and minerals are often redistributed throughout the entire plant. Indeed, we observed *BSCL1*, *BSCL2*, and *BSCL3* as being noticeably abundant in the roots that were comparable to that in the feeding sites and less than in the top leaves (Fig. 1*F*), when the feeding experiments were repeated with 14-day-old seedlings growing in hydroponics. In all experiments, the *BSCL*s were not detected in the control plants that were uninfested by BPH (Fig. S2 *D* and *E*). It strongly implies that the migration of *BSCL*s in rice plants is in alignment with the phloem streaming moving from the source to sink tissues.

### Visualization of *BSCL1* localization in planta

We examined the in planta migration of *BSCL1* using an RNA imaging approach, the RNA switch–controlled RNA-triggered fluorescence (RNA switch–RTF) system, which enables real-time visualization of RNA dynamics in living plants (20). In this system, the RNA switch with a probe targeting *BSCL1* can toggle between active and inactive states depending on the presence of the target RNA, thereby controlling GFP accumulation or degradation (Fig. 2*A*).

**Fig. 2.**
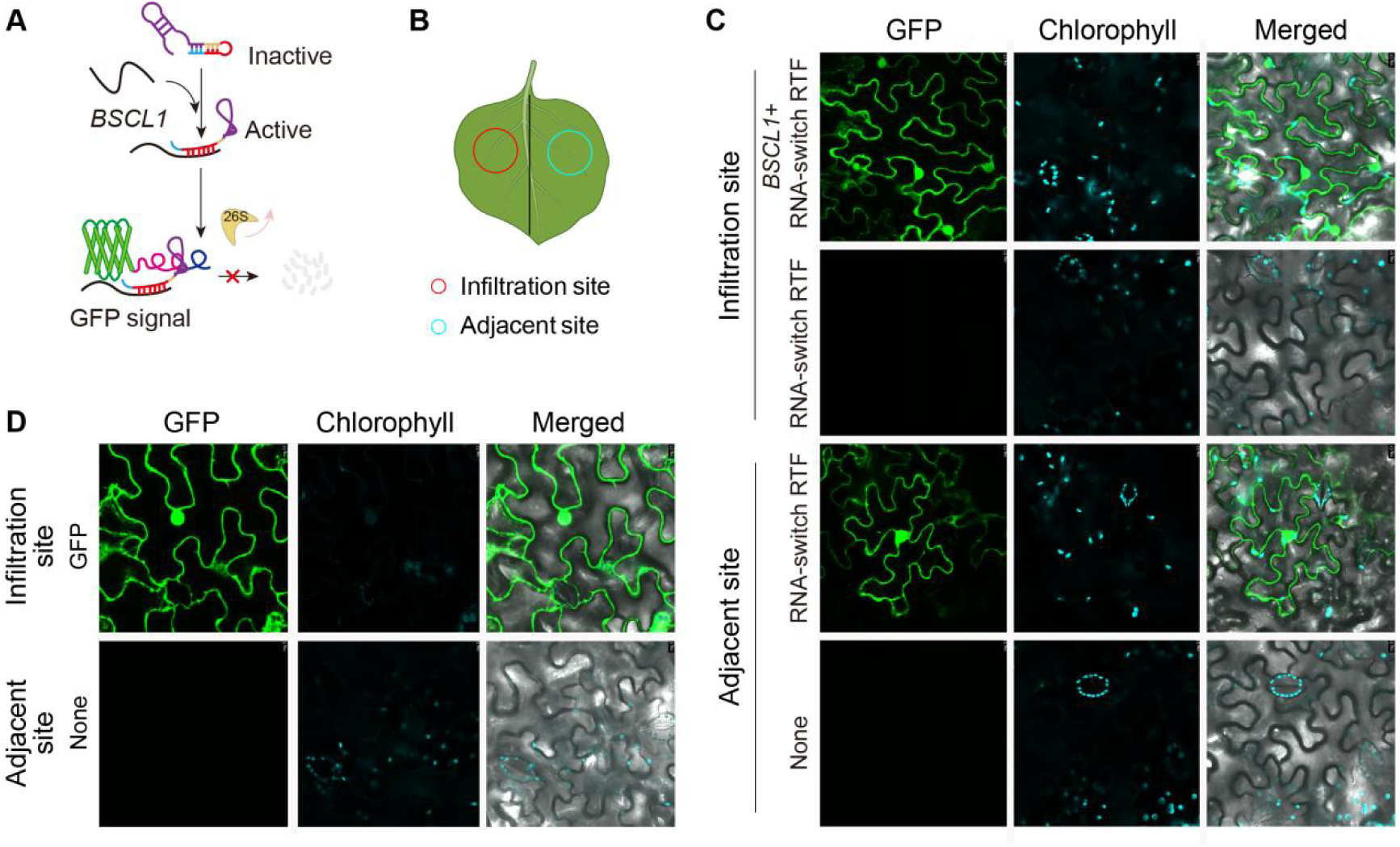
Visualization of *BSCL1* localization and movement. (A) Schematic of the RNA-switch–RTF reporter system. The red sequence in RNA-switch represents the probe for *BSCL1*. In the absence of target RNA, the RNA-switch remains inactive due to base pairing between the linkers and their complementary sequences in the probe, leading to GFP degradation via 26S proteasome. Upon target RNA binding, the probe hybridizes with the target, triggering the switch to an unfolding active state, recruiting in GFP accumulation at the RNA site (For details, please refer to the Materials and Methods section). (B) Schematic of infiltration injection site in *N. benthamiana* leaves. (C) Visualization of *BSCL1* localization and movement in *N. benthamiana* leaves using the RNA-switch RTF system. The infiltration site was injected with either a mixture of the *BSCL1* expression construct and the RNA-switch–RTF construct, or with the RNA-switch–RTF construct alone, while the adjacent site was treated with the RNA-switch–RTF construct accordingly. (D) GFP expressed alone is unable to move in *N. benthamiana* leaves. None indicated no construct infiltrated.

To design a specific probe, we first determined the full-length *BSCL1* sequence (708 nt) from brown planthopper (BPH) using 5′ and 3′ RACE (Fig. S4 *A* and *B*) and validated it by northern blotting (Fig. S4*C*). The full-length transcript was detected in FS, NFS, and DFS tissues (Fig. S4*D*). Based on this sequence, we designed a dumbbell-shaped RNA switch with the probe (Fig. S4*E*), which is essential for the RNA switch function.

Agroinfiltration of *N. benthamiana* leaves with *35S::BSCL1* together with the RNA switch–RTF construct produced GFP signals in both the nucleus and cytoplasm (Fig. 2 *B* and *C*), confirming subcellular localization of *BSCL1* (Fig. 2*C*). No GFP fluorescence was observed when leaves were infiltrated with either *BSCL1* or the RNA switch–RTF system alone (Fig. 2*C*). Remarkably, GFP signals were also detected in tissues adjacent to the infiltration sites, where the RNA switch–RTF system had been introduced but not the *BSCL1* plasmid (Fig. 2*C*), demonstrating systemic movement of *BSCL1*. By contrast, infiltration with *35S::GFP* produced strong local fluorescence but no signals in neighboring tissues (Fig. 2*D*).

Together, these results demonstrate that BPH-derived *BSCL*s are not only localized within plant cells but also migrate systemically from the initial delivery sites to adjacent and distal tissues.

### BPH *BSCLs* are more stable than BPH mRNAs in rice plants

A substantial number of BPH mRNA reads were also found in the RNA-seq of FS (Fig. S5*A*, Table S2). This prompted us to investigate whether BPH mRNAs also migrated in the rice plants. Out of seven selected BPH mRNAs (Fig. S5*A*), two (*Nlug010593* and *Nlug002054*) were found in the FS by PCR (Fig. 3*A*, Fig. S5*B*).

**Fig. 3.**
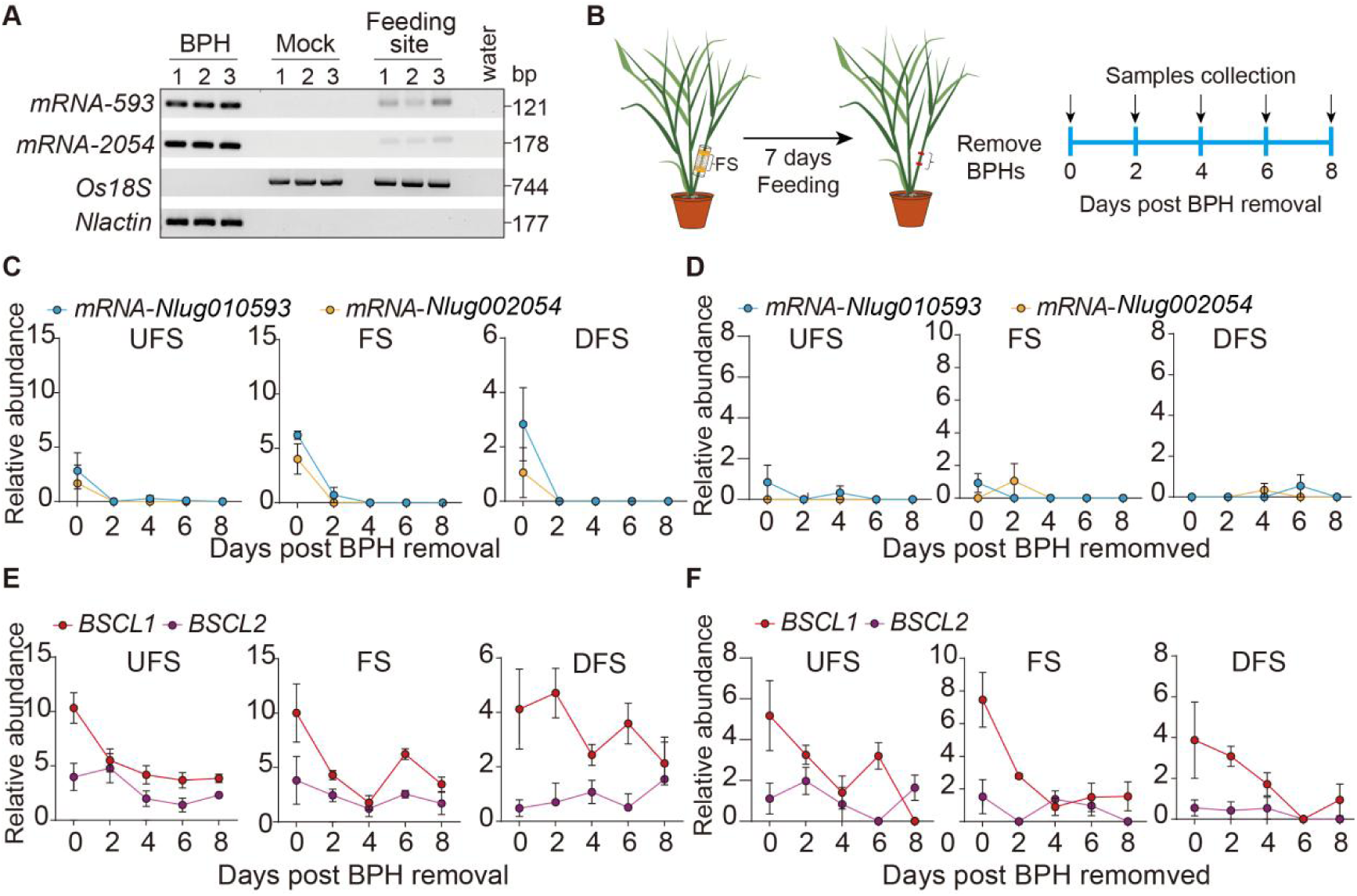
*BSCL*s persist in rice plants whereas BPH mRNAs are rapidly degraded. (A) BPH mRNAs were translocated into the FS of rice plants. (B) Schematic of sampling strategy showing collection at different time points post-BPH infestation. (C, D) The relative abundance of *mRNA-Nlug010593* and *mRNA-Nlug002054* transcripts declined rapidly in both the outer (C) and inner (D) layers of UFS, FS, and DFS across time points. (E, F) In contrast, *BSCL1* and *BSCL2* transcripts persisted for extended periods in both the outer (E) and inner (F) layers of UFS, FS, and DFS. A schematic of the inner and outer layers is provided in Fig. 2D. Data are shown as mean ± SE (n = 4 biological replicates).

However, the presence of these mRNAs in rice plants can only be detected by primers targeting the shorter fragments (Fig. S5 *C* and *D*), indicating that BPH mRNAs in the rice plants have been fragmented. Fragmented RNAs are often less stable and more prone to degradation (21, 22). To assess the stability of BPH mRNAs and *BSCL*s in rice plants, we removed BPH from FS after 7 days of feeding and subsequently harvested samples at several time points (Fig. 3*B*). On the 7^th^ day of feeding (0 days post-BPH removal), *BSCL*s (*BSCL1* and *BSCL2*) and BPH mRNAs (*Nlug010593* and *Nlug002054*) were found in both the outer and inner layers of FS (Fig. 3 *C* and *D*). The abundance of *BSCL1* and *BSCL2* remained relatively higher than that of *Nlug010593* and *Nlug002054* (Fig. 3 *C*-*F*). Two days posted BPH removal, the abundance of BPH mRNAs in the rice plants declined sharply. In contrast, the abundance of *BSCL*s in the rice plants remained high, even 8 days after the removal of BPH (Fig. 3 *C*-*F*), suggesting BPH *BSCL*s are more stable in rice plants than BPH mRNAs.

### BPH *BSCL1* is a virulence factor that suppresses the plant defenses

To investigate whether *BSCL*s contributed to the BPH colonization on rice plants, we selected *BSCL1* to conduct further experiments since *BSCL1* is the most abundant in FS compared to other *BSCL*s. We knocked down the expression levels of *BSCL1* in BPH through two different dsRNA sequences. The results showed that we had successfully inhibited the expression levels of *BSCL1* in BPH (Fig. 4*A*). The survival rate had no significant difference between *dsBSCL1* and the *dsGFP* groups (Fig. 4*B*), however, the honeydew production (Fig. 4*C*, Fig. S6*A*) and the total number of eggs per female (Fig. 4*D*) were significantly reduced.

**Fig. 4.**
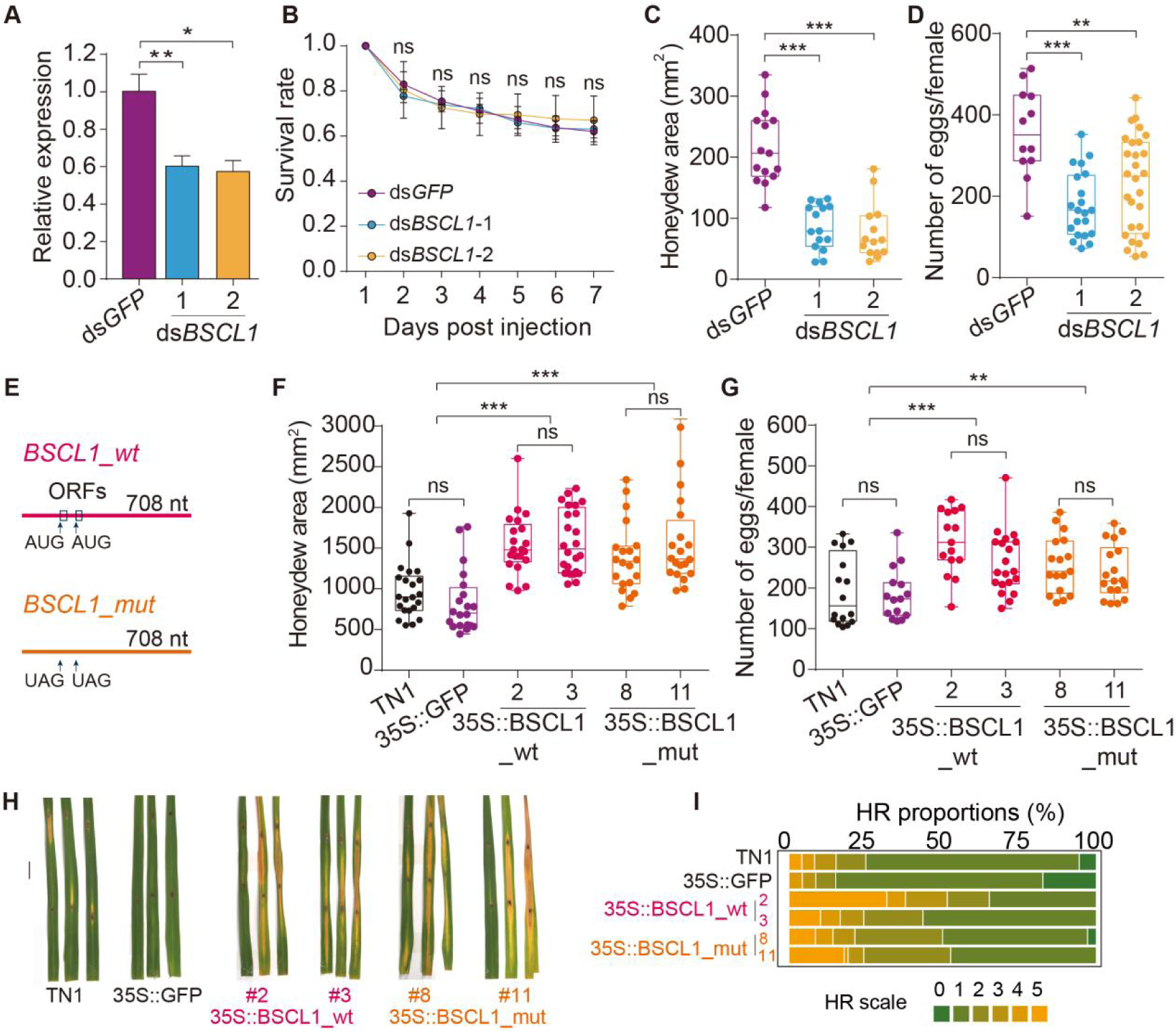
*BSCL1* promotes BPH colonization by suppressing rice defense responses. (A) Silencing of *BSCL1* by dsRNA injection. *dsGFP* served as a negative control. Data are means ± SE from three independent biological replicates. (B) Survival rates were unaffected by dsRNA treatment (90–100 insects per replicate, three replicates per treatment). (C, D) Honeydew excretion (C) and fecundity (D) were reduced after *BSCL1* silencing (n = 15 and n = 12–28 biological replicates, respectively). (E) Schematic of transgenic rice construction. (F, G) Honeydew excretion (F) and fecundity (G) were increased in BPH feeding on rice stably expressing *BSCL1_wt* or *BSCL1_mut* (n = 20–24 and n = 15–20 biological replicates, respectively). (H, I) Transgenic expression of *BSCL1_wt* or *BSCL1_mut* increased rice susceptibility to *M. oryzae*. (H) Disease symptoms caused by *M. oryzae* strain Guy11 at 7 dpi. (I) Hypersensitive response proportion of each rice line in (H). Stacked bars are color-coded to show the proportions (in percentage) of each hypersensitive response scale (0–5) out of total infiltrated leaves scored (For details, please refer to the Materials and Methods section). Total of 40 leaves are scored for each stacked column. Statistical significance was determined by paired Student’s *t* test: \*\*\**P* < 0.001, \*\**P* < 0.01, \**P* < 0.05, ns = not significant.

To further assess the impact of *BSCL1* on BPH performance, we generated stable transgenic plants in TN1 genetic background that produced 708-nt of *BSCL1* (35S::BSCL1, lines 2 and 3) and 708-nt *BSCL1* mutants in which two ATG start sites were mutated to stop codes (35S::BSCL1_mut, lines 8 and 11) (Fig. 4*E*). BPH produces more honeydew (Fig. 4*F*, Fig. S6*B*) and eggs (Fig. 4*G*) on both 35S::BSCL1 and 35S::BSCL1_mut compared to 35S::GFP and TN1 plants, suggesting *BSCL1* in rice promote BPH colonization. These findings suggest that BPH *BSCL1* is a virulence factor that plays a critical role in BPH-rice interactions.

To assess whether plant defense pathways were affected by *BSCL1*, we challenged TN1, 35S::GF plants, 35S::BSCL1_wt, and 35S::BSCL1_mut by *M. oryzae*. Six days posted the inoculation of *M. oryzae*, 35S::BSCL1_wt and 35S::BSCL1_mut plants exhibited a stronger hypersensitive response compared to TN1 and 35S::GFP plants (Fig. 4 *H* and *I*), suggesting that overall rice defense pathways were suppressed by *BSCL1*.

### *BSCL1* represses the expression of defense-related genes in rice

To determine the host processes influenced by *BSCL1*, we performed RNA-seq on rice plants (TN1, 35S::GFP, 35S::BSCL1_wt, and 35S::BSCL1_mut) with or without BPH feeding for 24 h. Principal component analysis (PCA) of transcriptomes revealed clear separation of sample groups (Fig. S7*A*). Relative to non-feeding controls, BPH feeding altered the expression of 6,710 rice genes (Fig. S7*B*, Dataset S4).

We next examined whether *BSCL1* modulates these plant responses. Coexpression analysis of differentially expressed (DE) genes grouped 6,404 genes into eight clusters based on expression patterns (Fig. 5*A*; Dataset S5). TPM heatmaps confirmed these clusters (Fig. S8). Expression profiles of 35S::BSCL1_mut plants closely resembled those of 35S::BSCL1_wt, indicating that *BSCL1* functions as an RNA rather than through a peptide product (Fig. S8).

**Fig. 5.**
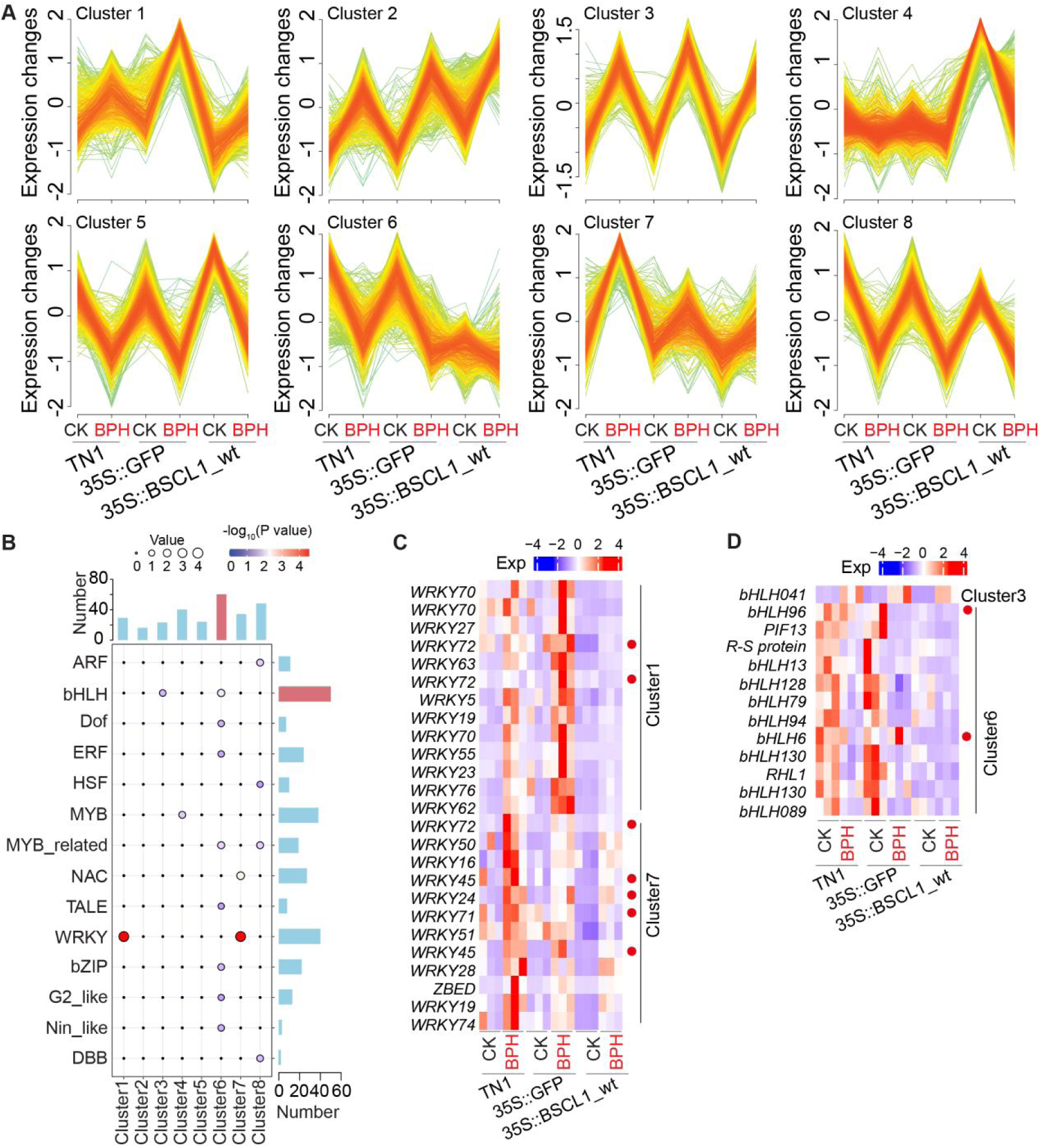
*BSCL1*-induced DE genes are enriched for defense-related TFs. (A) Expression trends of DE genes grouped into eight clusters. CK, uninfested plants; BPH, plants infested by BPH. The colors indicate the membership belonging to the clusters. Red, yellowish, and greenish rank from high, medium, and low. (B) Enrichment of TFs across clusters. Bar plots on the right and top indicate the total number of TFs per family and per cluster, respectively. Red marker bars indicate the highest values in each plots. (C, D) Heatmaps of TPM for enriched *WRKY* genes in clusters 1 and 7 (C) and *bHLH* gene families in clusters 3 and 6 (D). Genes marked with red dots have been previously reported to be associated with rice defense against BPH [*WRKY72* (37), *WRKY24* (38), *WRKY71* (39), *WRKY45* (40), *bHLH96* (23), *bHLH6* (24)].

Four clusters (clusters 2, 3, 5, and 8) showed similar expression in 35S::BSCL1_wt, TN1, and 35S::GFP plants, suggesting that these groups are largely unrelated to *BSCL1* activity (Fig. 5*A*; Fig. S8 *B*, *C*, *E*, and *H*). Cluster 4 genes were strongly induced in 35S::BSCL1_wt plants but suppressed after BPH feeding, a pattern absent in controls (Fig. 5*A*; Fig. S8*D*). By contrast, clusters 1, 6, and 7 exhibited distinct expression changes in 35S::BSCL1_wt plants (Fig. 5*A*; Fig. S8 *A*, *F*, and *G*). In clusters 1 and 7, many genes were strongly upregulated in TN1 and 35S::GFP plants after BPH feeding, but this induction was markedly repressed in 35S::BSCL1_wt plants (Fig. 5*A*; Fig. S8 *A* and *G*). These clusters were significantly enriched for genes in the jasmonic acid (JA)-mediated defense pathway (Fig. S9). Consistently, multiple JA pathway genes were repressed in both 35S::BSCL1_wt and 35S::BSCL1_mut compared to controls (Fig. S10).

Cluster 6 was of particular interest because its genes were downregulated upon BPH feeding and remained further suppressed in 35S::BSCL1_wt plants (Fig. 5 *A* and *B*). Notably, cluster 6 contained the highest number of transcription factor (TF) genes among all clusters (Fig. 5*B*). Enriched families included bHLH, MYB-related, bZIP, Dof, Nin-like, ERF, G2-like, and TALE, with bHLH TFs most abundant (12/38; Fig. 5*B*). Several members of this family—including bHLH96, bHLH6, and PTF1—are known regulators of JA- and SA-mediated defenses. Rice bHLH96 and bHLH6, in particular, have been implicated in resistance to BPH (23, 24). Their repression in both control plants and 35S::BSCL1_wt upon BPH feeding suggests that *BSCL1* interferes with transcriptional regulation of key defense pathways (Fig. 5 *C* and *D*, Table S3).

### *BSCL1* targets the histone complex to repress transcription factor promoters

In plants, lncRNAs have been reported to act through protein interactions (25, 26). To identify *BSCL1*-binding partners in rice, we performed yeast three-hybrid (Y3H) screening using a rice cDNA library. A total of 197 yeast colonies grew on selective media, of which 104 exhibited strong X-β-gal activity (Fig. S11, Dataset S6). These corresponded to 31 rice proteins, including enzymes, transcription factors, and components of the histone complex (Table S4). Notably, vacuolar protein sorting-associated protein 2 homolog 2 (VPS), histone H3.3, transcription factor RF2a-like (RF2), ribonuclease J isoform X1, HIRA-interacting protein 3 (HIRIP3), and T-complex protein 1 subunit gamma were among the frequently recovered candidates.

Given the central role of chromatin regulation in transcriptional control, we focused on histone H3.3 and HIRIP3. Interactions of *BSCL1* with both proteins were validated in repeated Y3H assays (Fig. 6*A*). RNA-switch-RTF revealed that *BSCL1* was colocalized with HIRIP3 in the nucleus and cytoplasm, with H3.3 in the nucleus, where H3.3 was mainly expressed (Fig. 6*B*). These results suggest that *BSCL1* associates with the HIRA histone chaperone complex, which normally deposits H3.3 into chromatin to maintain active transcription states.

**Fig. 6.**
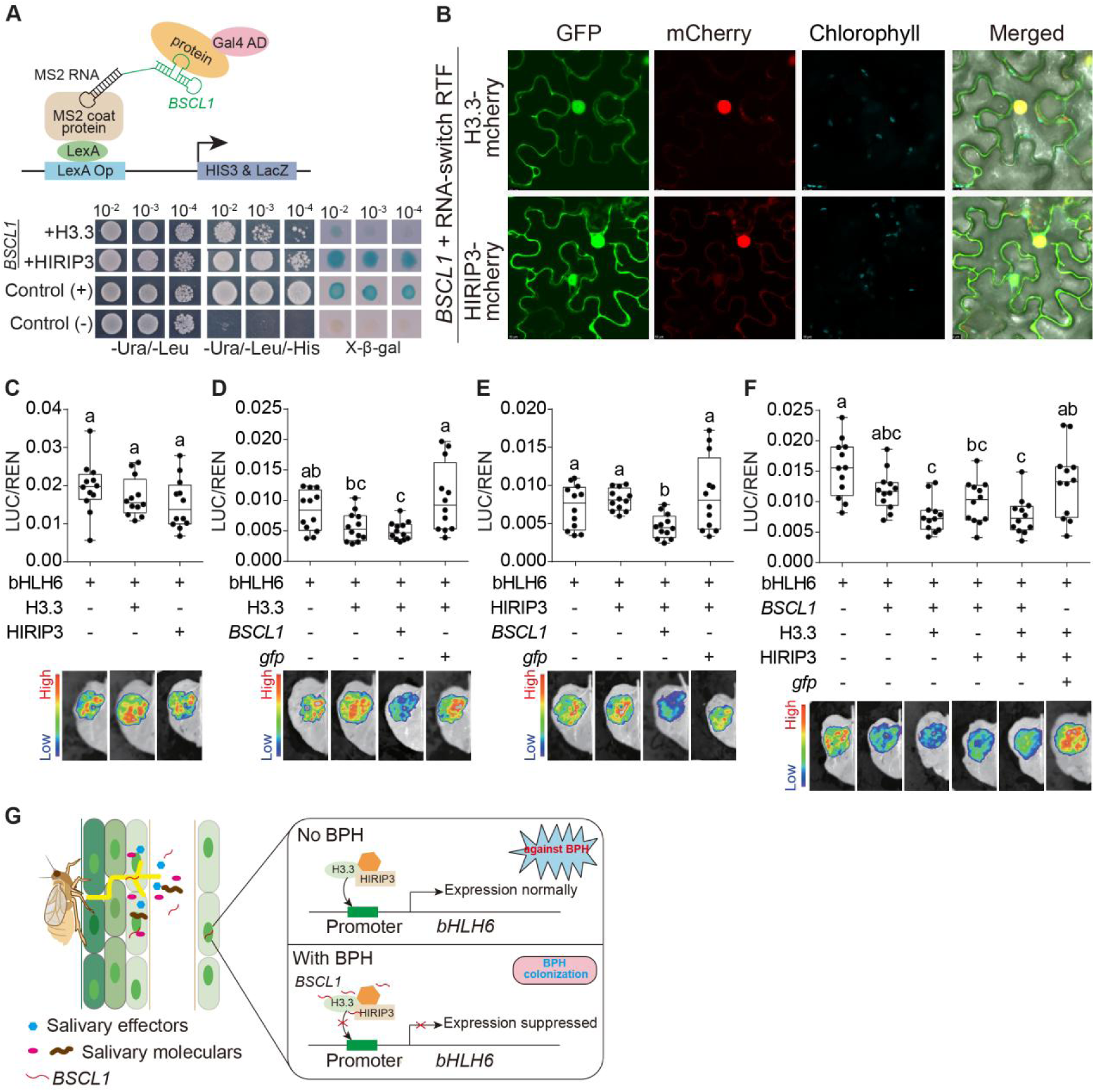
*BSCL1* evicts the histone complex from the *bHLH6* promoter and represses its transcription. (A) Yeast three-hybrid screening identified H3.3 and HIRIP3 as *BSCL1*-interacting proteins. MS2 sequences fused to *BSCL1* served as bait, and a rice cDNA library fused to the Gal4 activation domain served as prey. Interaction activated reporter genes (*HIS3* and *lacZ*). (B) Co-localization of *BSCL1* with H3.3 and HIRIP3 in the nucleus and cytoplasm of *N. benthamiana* epidermal cells. *BSCL1* localization was visualized with the RNA-switch–RTF system, and H3.3/HIRIP3 were fused to mCherry. (C) Expression of H3.3 or HIRIP3 alone did not affect luciferase activity driven by the *bHLH6* promoter. (D–F) Co-expression of *BSCL1* with H3.3 (D) or HIRIP3 (E), or both (F) suppressed *bHLH6* promoter activity. Boxplots show luciferase activity (n = 12). Bioluminescence images of representative treatments are shown below. Different lowercase letters indicate significant differences (*P* < 0.05, one-way ANOVA with Tukey’s test). (G) Proposed model of *BSCL1* function. In uninfested plants, the histone complex regulates *bHLH6* transcription, contributing to rice resistance to BPH. During infestation, *BSCL1* is secreted into rice, where it disrupts histone–DNA interactions at the *bHLH6* promoter, repressing transcription and promoting BPH colonization.

We next tested whether *BSCL1* interferes with H3.3 deposition at promoters of defense-related transcription factors (e.g., bHLH6 in cluster 6). A luciferase (LUC) reporter driven by the bHLH6 promoter was co-expressed with combinations of H3.3, HIRIP3, and *BSCL1* in rice. Neither H3.3 nor HIRIP3 alone altered promoter activity (Fig. 6*C*), but co-expression with *BSCL1* significantly suppressed LUC expression (Fig. 6 *D* and *E*). Importantly, simultaneous expression of H3.3 and HIRIP3 did not produce further repression beyond that observed with either factor alone (Fig. 6*F*), indicating that *BSCL1* disrupts the function of the H3.3–HIRIP3 complex at defense gene promoters (Fig. 6*G*).

## Discussion

We found that BPH actively translocated salivary gland RNAs into rice plants during feeding. These cross-kingdom RNAs move systemically from feeding sites on the sheath to distant tissues such as inner sheaths, upper leaves, and roots, likely through vascular transport. In rice, BPH mRNAs degraded relatively quickly—possibly via 5′-end decapping and processing as part of a conserved host defense strategy—whereas BPH lncRNAs persisted for extended periods, remaining detectable up to eight days after insect removal. Among these lncRNAs, *BSCL1* emerged as a highly abundant and functionally important effector. Silencing *BSCL1* in BPH reduced feeding and reproduction, while heterologous expression in rice enhanced BPH performance. Moreover, *BSCL1* overexpression suppressed JA-induced defense transcription factors and compromised immunity against both BPH and the blast fungus *M. oryzae*, suggesting that it broadly disables rice plant defense.

Transcriptomic profiling revealed that BPH feeding orchestrates complex reprogramming of rice metabolism and defense. We observed repression of processes associated with cell walls, ion transport, and carbohydrate metabolism, but strong induction of jasmonic acid (JA)-related genes, RNA metabolism, circadian rhythm, and transporter activities (Fig. S8). These patterns are consistent with previous findings: reinforcement of the cell wall enhances resistance (27–29); ion transport underpins defense signaling and metabolite biosynthesis (17); and carbohydrate metabolism, often manipulated by herbivores, is further exploited by BPH through hijacking SWEET13/14 sugar transporters (30–35). Despite these suppressions, BPH feeding activates defense signaling, with strong induction of JA pathway genes and transcription factors such as WRKY, NAC, and MYB (23, 36–40) and tissue-specific JA sectors including the MYC2–bHLH6 cascade (24). Together, these findings highlight the dual nature of the interaction: rice perceives and counters insect attack, while BPH simultaneously manipulates host physiology to promote feeding.

Our discovery of insect-derived lncRNAs reveals a previously unrecognized mechanism: BPH not only manipulates host physiology indirectly but also delivers its own genetic effectors to reprogram rice immunity at transcriptional and epigenetic levels. *BSCL1* exemplifies this new class of mobile effectors. It suppresses JA-associated transcription factors such as OsbHLH6—key regulators of defense (23, 24)—and interacts with the HIRA histone chaperone complex to block H3.3 deposition at their promoters (Fig. 6*G*). Similar lncRNA-guided epigenetic mechanisms have been described in other systems (41, 42). These findings also force a reinterpretation of prior observations. The downregulation of defense TFs reported (36) and the apparent negative feedback in JA signaling (24) may partly result from direct suppression by insect lncRNAs. Likewise, sugar transport manipulations by BPH (35) likely occur in a defense-compromised context where RNA effectors such as *BSCL1* have already silenced immune hubs. Thus, BPH deploys a multilayered strategy: altering host nutrition, modulating hormone signaling, and injecting RNA-based epigenetic suppressors that dismantle transcriptional control of immunity.

BPH secretes a cocktail of salivary molecules to manipulate rice and promote feeding, including enzymatic proteins (43), effector proteins (44, 45), and miRNAs (5). In addition, numerous salivary gland–derived lncRNAs and mRNAs are translocated into rice, potentially trafficking systemically via the phloem. BPH mRNAs are rapidly degraded, consistent with a conserved host defense against foreign RNAs, as reflected by enrichment of RNA metabolism pathways in Cluster 2 upon BPH feeding (Fig. S8). In contrast, a small subset of RNAs, including *BSCL1*, remains stable as full-length transcripts, suggesting structural features that allow evasion of host RNA surveillance. While cross-kingdom transfer of long RNAs has been reported in parasitic plants, fungi, and aphids—including the aphid lncRNA *Ya1* shown to function as an effector (6–8, 10)—*BSCL1* exemplifies these principles and raises key questions: can some insect mRNAs escape host surveillance and be translated in plant cells, as reported for fungal effectors (8)? How do cross-kingdom lncRNAs such as *BSCL1* and *Ya1* evade detection, and how do resistant versus susceptible rice genotypes influence their stability, systemic movement, and function? Answering these questions will provide critical insight into the molecular strategies insects use to manipulate host immunity and may guide innovative approaches for pest control.

In sum, the discovery of BPH *BSCLs* reveals lncRNAs as systemic epigenetic effectors that reprogram host immunity. This shifts the view of insect effectors from plant-centric to insect-centric and highlights lncRNAs as an overlooked class of cross-kingdom regulators. Beyond advancing basic understanding of plant–herbivore interactions, these findings suggest new strategies for pest control by targeting RNA effectors.

## Author contributions

Y.C. and H.H. conceived and designed the study, supervised the project, and finalized the manuscript. D.W. carried out the majority of the experimental work. S.J. and Z.Q. contributed to the experimental setup and bioassays. Bioinformatic analyses were conducted jointly by Y.C. and D.W. C.L., J.W., R.H., and Y.Z. contributed to the performed research. Y.H. and W.M. contributed to analyzed data and experiment design. The manuscript was drafted by Y.C. and D.W., with all authors contributing to its revision. All authors reviewed the manuscript, provided feedback, and approved the final version.

## Data and materials availability

All data reported in this paper were deposited in the NCBI BioProject under project numbers PRJNA1197768, PRJNA1314766, and PRJNA1330246. All data are available in the main text or the supplementary materials.

## Acknowledgments

This project is funded by the National Key Research and Development Program of China (project No. 2024YFD1400700 to H.H.), the Hubei Grain Yield Improvement Program (project No. 2662025YJ017 to H.H.), the National Key Research and Development Program of China (project No. 2023YFF1000703 to Y.C.), the National Natural Science Foundation of China (project No. 32172392 to Y.C.), the Hubei Hongshan Laboratory (project No. 2022hszd026 to Y.C.), the Startup Foundation for Advanced Talents at HZAU to Y.C., the Fundamental Research Funds for the Central Universities (Program No. 2022ZKPY003 to Y.C.), and the Wuhan Yingcai Talent Program to Y.C.

## Supplementary Information

### Supplementary text

#### Materials and methods

##### Insect rearing and plant growth

Brown planthopper (BPH, *N. lugens*) populations were originally collected from rice fields in Wuhan, China, and have been continuously maintained on the susceptible rice cultivar Taichung Native 1 (TN1) since 2007.

Rice materials used in this study included TN1, the resistant cultivar Rathu Heenati (RHT) (1), and transgenic lines (35S::GFP, 35S::BSCL1_wt, and 35S::BSCL1_mut). Rice seeds were sown in plastic pots (7.5 cm diameter, 10 cm height; one plant per pot) and placed in the fields of Huazhong Agricultural University. Plants were protected with insect-proof cages to prevent pest infestation. Tillering stage (30∼45 day-old) plants were brought back to the laboratory for subsequent assays, and were grown under the controlled conditions of 27 ± 1 °C, 70 ± 5% relative humidity, and a 14:10 h light/dark photoperiod.

Hydroponically grown TN1 seedlings were germinated in Petri dishes (9 cm diameter, 2 cm height) containing water and incubated in a controlled growth chamber under the same controlled conditions mentioned above.

##### Identification and evolutionary analysis of BPH lncRNAs

To identify and analyze BPH lncRNAs, we employed a computational pipeline integrating Evolinc-I for lncRNA discovery and Evolinc-II for evolutionary conservation analysis (2). RNA-seq datasets from BPH (NCBI project PRJNA514182) (3) were first quality-checked using FastQC (v0.11.5) (4) and low-quality bases reads were removed. RNA seq reads were mapped to the BPH genome using the RMTA (v2.6.3) pipeline (5) with default parameters. Transcript assembly was performed using StringTie (v2.2.3) (6), and assembled GTF files were merged with Cuffmerge (v2.2.1.5) (7) to generate a comprehensive transcriptome. The Evolinc-I pipeline was then applied to identify candidate lncRNAs. Transcripts overlapping annotated protein-coding genes or transposable elements, or shorter than 200 nucleotides, were excluded. Coding potential was evaluated using CPC2 (v2.0) (8), and transcripts with significant coding potential were removed. Remaining transcripts were further screened against the Rfam database (v15.0) (9) to exclude known structural RNAs, including tRNAs, rRNAs, and snoRNAs. The resulting set of transcripts was considered putative lncRNAs.

To assess the evolutionary conservation of BPH lncRNAs, we utilized the Evolinc-II pipeline. This tool performs reciprocal BLAST analyses (Evalue cutoff, 1e-20) against 28 arthropod genomes (Table S5). Homologous sequences were grouped into families based on sequence similarity.

##### BPH feeding on different resistant rice lines

At the tillering stage (∼30 days), TN1 (susceptible) and RHT (resistant) rice plants were each infested with 30 third-instar BPH nymphs. After 48 h of feeding, surviving nymphs were collected and immediately snap-frozen in liquid nitrogen. Three biological replicates were performed per cultivar, and all samples were stored at -80 °C for RNA extraction.

##### BPH feeding experiments

To test whether BPH secreted transcripts into rice, 30 male adults, to avoid egg deposition by females, were confined on stems of 45-day-old TN1 plants (tillering stage) using glass cylinders (10 cm length, 2.5 cm diameter). Uninfested plant stems with empty glass cylinders served as controls. After 48 h of feeding, feeding sites (FS) were excised, rinsed three times with deionized water, and snap-frozen in liquid nitrogen. Each treatment was conducted with two to three biological replicates. The experiment was repeated three times, achieving three replicates for controls and seven replicates for the FS. Samples were used for RNA-seq to identify BPH transcripts potentially translocated in the rice plants.

##### RNA extraction, library preparation, and sequencing

RNA-seq was performed by OE Biotech Co., Ltd. (Shanghai, China). Samples were frozen in liquid nitrogen, ground to a fine powder using sterilized stainless-steel beads in a TissueLyser II (Jingxin, Shanghai, China), and homogenized in TRIzol reagent (Invitrogen, 15596018CN, USA) following the manufacturer’s protocol. RNA concentration and purity were measured with a NanoDrop 2000 spectrophotometer (Thermo Scientific, USA), and RNA integrity was assessed with an Agilent 2100 Bioanalyzer (Agilent Technologies, USA). Quantification was performed using a Qubit 2.0 Fluorometer (Thermo Scientific, USA). Only samples with RNA integrity number (RIN) >7.0 were used for library construction.

For each sample, 1 μg of high-quality total RNA was used to prepare strand-specific libraries with the VAHTS^®^ Universal V6 RNA-seq Library Prep Kit (Vazyme, NR604, Nanjing, China). The libraries were assessed for quality and fragment size distribution using the Bioanalyzer 2100. Sequencing was performed on an NovaSeq 6000 platform (Illumina, USA), generating 150-bp paired-end reads (project No. PRJNA1314766, PRJNA1197768, and PRJNA1330246).

##### RNA-seq analysis

RNA-seq datasets were analyzed from two sources: (i) publicly available data (PRJNA714229) covering diverse BPH tissues (antenna, fat body, gut, head, integument, ovipositor, ovary, and salivary gland) to investigate tissue-specific expression, and (ii) RNA-seq data generated in this study from BPH feeding on susceptible (TN1) and resistant (RHT) rice varieties to assess transcriptional responses to host resistance. The BPH reference genome (10) was used for read alignment, while feeding site samples were aligned against a merged reference of TN1 rice (11) and BPH (10).

Raw sequencing reads were processed with fastp (v0.20.1) (12) to remove adapter sequences and low-quality reads (phred score <25). Reads shorter than 50 bp after trimming were discarded. The quality of clean reads was evaluated using FastQC (v0.11.5) (4), and only reads passing quality control were retained for downstream analysis.

Clean reads were aligned to the reference genome using RMTA (v2.6.3) (13) with the HISAT2 aligner. Parameters were set to trim 15 bases from the 5′ end of each read, and minimum and maximum intron lengths were specified as 20 and 500,000, respectively. The resulting BAM files were used for read quantification with HTSeq (v0.6.1) (14), using the parameters -r, -i gene_id, -t exon, and library-specific strand settings (-s no for non-stranded or -s yes for stranded data). Only uniquely mapped reads overlapping annotated exons were counted.

Differential gene expression was analyzed with edgeR (v3.0) (15). Raw read counts from HTSeq were filtered to remove lowly expressed genes (counts per million, CPM <1), normalized using the trimmed mean of M values (TMM) method, and tested for differential expression. P-values were corrected for multiple testing using the Benjamini–Hochberg method. Genes with false discovery rate (FDR)-adjusted *p* < 0.05 and |log2 fold change| ≥ 1 were considered significantly differentially expressed.

Transcript abundance was also quantified as TPM (transcripts per million) using TPMCalculator (v0.0.3) (16). BAM files generated by RMTA and the corresponding GTF annotation files were provided to TPMCalculator (-b for BAM, -g for annotation). Strand-specificity was set according to the RNA library type (--stranded no for non-stranded libraries).

##### Migration of BPH-derived RNAs in rice plants

To examine the systemic movement of BPH-delivered RNAs in rice, feeding experiments were performed at different developmental stages of TN1 plants. For tillering-stage plants (∼30-day old), 50 fourth-instar BPH nymphs were confined on a stem using a glass cylinder cage (10 cm length, 2.5 cm diameter) and allowed to feed for 48 h. After feeding, tissues were collected from two additional positions relative to the feeding site (FS): 2 cm above the FS (UFS), 2 cm below the FS (DFS).

To better resolve the spatial distribution of transcripts within stem tissues, FS, UFS, and DFS samples were dissected into two fractions: (i) outer layers (the two outermost sheaths) and (ii) inner layers (remaining tissues). All collected tissues were rinsed three times with sterile deionized water to remove surface contamination and immediately snap-frozen in liquid nitrogen. Control plants were treated with empty cages. Each treatment included three biological replicates, and the entire experiment was repeated three independent times.

For hydroponic experiments, hydroponically grown TN1 plants at the three-leaf stage (∼14-day old) were used. 20 fourth-instar nymphs were confined on the stems of four seedlings per cage for 48 h. After feeding, tissues were harvested from the leaf, FS, DFS, and root. Each tissue sample was thoroughly washed three times with sterile deionized water prior to freezing in liquid nitrogen. Seedlings caged without insects were used as controls. Each treatment was performed with three biological replicates, and the experiment was repeated three independent times.

##### Experiments to analyze BPH RNA stability in rice plants

To assess the stability of BPH-delivered mRNAs and lncRNAs in rice, a time-course feeding experiment was performed. Approximately 80 third-instar BPH nymphs were caged on stems of TN1 rice at the tillering stage (∼30-day old) using glass cylinders (10 cm length, 2.5 cm diameter) and allowed to feed for seven days. After feeding, all insects were removed, and plant tissues were harvested at 0, 2, 4, 6, and 8 days post-removal. Four biological replicates were collected per time point. Tissues were dissected into outer and inner sheath layers, thoroughly washed with sterile deionized water, and snap-frozen in liquid nitrogen.

The relative abundance of cross-kingdom RNAs was quantified by qRT-PCR. Two mRNAs (*Nlug010593*, *Nlug002054)* and two lncRNAs (*BSCL1*, *BSCL2*) were examined at multiple tissue sites (FS, UFS, DFS) across all time points.

##### RNA extraction, cDNA synthesis, qRT-PCR

Total RNA was extracted from collected tissues using TRIzol reagent (Invitrogen, 15596018CN, USA) and treated with RNase-free DNase I (Thermo Fisher Scientific, EN0521, USA) to remove residual genomic DNA. First-strand cDNA was synthesized from 1 μg of total RNA using a mixture of oligo (dT) and random primers with the RevertAid First Strand cDNA Synthesis Kit (Thermo Fisher Scientific, K1622, USA), following the manufacturer’s instructions.

Relative transcript abundance was measured by qRT-PCR. Two BPH mRNAs (*Nlug010593*, *Nlug002054*) and two lncRNAs (*BSCL1*, *BSCL2*) were analyzed at multiple tissue sites (FS, DFS, and UFS) across all time points. qRT-PCR reactions were performed in 20 μL volumes containing 10 μL SYBR Green (Takara, RR820A, Japan), 0.5 μL of each primer (10 μM), 1 μL of cDNA template, and 8 μL nuclease-free water, using the CFX Connect™ Real-Time System (Bio-Rad, USA). Cycling conditions were 95°C for 10 s, followed by 40 cycles of 95°C for 20 s, 60°C for 30 s, and 72°C for 30 s. Expression levels were normalized to *O. sativa* 18S rRNA (*Os18S*) and quantified using the 2^^-ΔCt^ method. Primer sequences are listed in Table S6.

##### Cloning and sequencing of *BSCL1*

To obtain the full-length cDNA of *BSCL1*, 5’ and 3’ rapid amplification of cDNA ends (RACE) was performed. For 3’ RACE, 3 μg of BPH total RNA was added to an 80 μL ligation mixture containing 4 μL T4 RNA ligase, 8 μL T4 RNA ligase buffer, 8 μL BSA (Thermo Fisher Scientific, EL0021, USA), 8 μL ATP (Thermo Fisher Scientific, R0481, USA), and 10 pmol of 3’ RACE RNA adaptor (Table S6). Ligation was carried out overnight at 16°C. The ligated RNA was reverse-transcribed into cDNA using an oligonucleotide complementary to the 3’ RACE adaptor with the RevertAid First Strand cDNA Synthesis Kit (Thermo Fisher Scientific, K1622, USA). For 5’ RACE, 4.1 μL of BPH total RNA was mixed with 10 μL of ligation mixture containing 0.1 μL T4 RNA ligase, 1 μL T4 RNA ligase buffer, 1 μL BSA (Thermo Fisher Scientific, EL0021, USA), 1 μL ATP (Thermo Fisher Scientific, R0481, USA), and 20 pmol 5’ RACE RNA adaptor (Table S6), followed by overnight ligation at 16°C. The ligated RNA was reverse-transcribed into cDNA using primers complementary to the 5’ RACE adaptor.

RACE PCRs were performed with *BSCL1* gene-specific forward or reverse primers together with the 3’ or 5’ RACE adaptor primers (Table S6). Each 50 μL PCR reaction contained 1 μL Phanta® Max Super-Fidelity DNA Polymerase (Vazyme, P505, Nanjing, China), 25 μL Phanta Max Buffer, 1 μL dNTP mix, 1 μL cDNA template, 1 μL of each primer (10 μM), and 20 μL nuclease-free water. The cycling program was: 95°C for 3 min, followed by 36 cycles of 95°C for 15 s, 55°C for 15 s, and 72°C for 20 s, with a final extension at 72°C for 5 min. PCR products were purified, ligated into the pEASY®-Blunt Cloning vector (TransGen, CB101, Beijing, China), and sequenced to confirm the full-length *BSCL1* cDNA.

##### Northern blotting

A 708-nt DNA fragment of *BSCL1* was cloned under a T7 promoter. *BSCL1* RNA transcript was synthesized using MAXIscript™ T7 Transcription Kit (Thermo Fisher Scientific, AM1314, USA). A Biotin-11-UTP (Thermo Fisher Scientific, AM8450, USA) labelled full-length of *BSCL1* antisense was synthesized using the MAXIscript™ T7 Transcription Kit (Thermo Fisher Scientific, AM1314, USA). 10 ng of *BSCL1* RNA and total BPH RNA were, respectively, mixed with RNA loading buffer (Takara, 9168, Japan), heated at 65 °C for 10 min, and separated on a 10% TBE-Urea denaturing polyacrylamide gel (Beyotime, R0235S, Shanghai, China) in 1 × TBE buffer at 150 V for 90 min. RNAs were transferred to a nylon membrane (Beyotime, FFN10, Shanghai, China) by electroblotting at 60 V for 60 min in 0.5 × TBE buffer and cross-linked to the membrane using a UV crosslinker (Ultraviolet Products, CA, USA): twice on the gel-facing side and once on the opposite side. The membrane was pre-hybridized in 10 mL hybridization solution at 42 °C for 1 h. The biotin-labeled *BSCL1* antisense was denatured at 70 °C for 10 min, cooled on ice for 5 min, and added to the hybridization solution for hybridization at 42 °C overnight.

Detection was carried out using the Chemiluminescent Nucleic Acid Detection Module Kit (Thermo Fisher Scientific, 89880, USA) following the manufacturer’s instructions. The membrane was blocked with 16 mL of Blocking Buffer for 15 min with gentle shaking. After decanting, 16 mL of conjugate/blocking solution was added, and the membrane was incubated for another 15 min. The membrane was washed four times for 5 min each with 20 mL 1 × wash solution under gentle shaking, then equilibrated in 30 mL Substrate Equilibration Buffer for 5 min. Chemiluminescent detection was performed by incubating the membrane in equal volumes of Luminol/Enhancer Solution and Stable Peroxide Solution for 5 min, followed by visualization using a CCD-equipped imaging system.

##### Generating overexpression rice plants

The *BSCL1* sequence is 708 nt in length (*BSCL1_wt*) and contains two AUG codons at positions 164 and 253. To generate a mutant form, both AUGs were replaced with UAGs using overlapping PCR (17), resulting in the *BSCL1_mut* sequence. For this, three overlapping fragments were amplified separately with primer pairs PC-*BSCL1*-F–*BSCL1*-ATG-164R (fragment 1), *BSCL1*-ATG-164F–*BSCL1*-ATG-253R (fragment 2), and *BSCL1*-ATG-253F–PC-*BSCL1*-R (fragment 3). Fragments 1 and 2 were first combined by overlap PCR with primers PC-*BSCL1*-F and *BSCL1*-ATG-253R. The resulting product was then fused with fragment 3 using primers PC-*BSCL1*-F and PC-*BSCL1*-R to obtain the 708-nt *BSCL1_mut* sequence. Primers were listed in the Table S6.

For rice transformation, the *BSCL1_wt*, *BSCL1_mut* and *GFP* sequences were cloned into the binary vector PC1300S under the control of the cauliflower mosaic virus 35S promoter. *BSCL1_wt* was amplified from BPH cDNA using primers carrying KpnI and XbaI restriction sites in the forward and reverse primers, respectively. *BSCL1_wt*, *BSCL1_mut*, and *GFP* were inserted into the PC1300S vector by homologous recombination using the ClonExpress Ultra One Step Cloning Kit (Vazyme, C115, Nanjing, China), yielding constructs PC1300S_35S::BSCL1_wt, PC1300S_35S::BSCL1_mut, PC1300S_35S::GFP. Plasmid constructs were first introduced into *Agrobacterium tumefaciens* strain EHA105 and verified by selection and molecular confirmation.

The generation of transgenic rice was performed according to the method described by Hiei and Komari (18), with minor modifications. Embryogenic calli were induced from surface-sterilized TN1 rice seeds and pre-cultured to increase transformation efficiency. The calli were then infected with the transformed *Agrobacterium* suspension and co-cultivated on medium containing acetosyringone to facilitate T-DNA transfer. Following co-cultivation, the calli were washed and placed on selective medium (N6 salts, N6 vitamins, 2,4-D 2 mg/L, Sucrose 30 g/L, Casein hydrolysate 0.3 g/L, Proline 0.5 g/L, Agar 8 g/L, Hygromycin B 50 mg/L, Cefotaxime 250 mg/L) containing the appropriate antibiotic to suppress bacterial growth and select for resistant transformants. Resistant calli were subsequently regenerated on hormone-supplemented medium (MS Basal Salt Mixture, MS Vitamin Stock, Sucrose 30 g/L, 6-BAP: 3.0 mg/L, NAA: 0.2 mg/L, Casein Hydrolysate 300 mg/L, Timentin 200 mg/L, Hygromycin 40 mg/L, Phytagel 3.2 g/L) to induce shoots, transferred to rooting medium (MS Basal Salt Mixture, MS Vitamin Stock, Sucrose 20 g/L, IBA 1.5 mg/L, Agar 7.0 g/L), and finally acclimatized in soil to obtain transgenic plants. Positive transformants were confirmed by PCR. The obtained positive plants of T2 generations were used for subsequent experiments.

##### Rice blast fungal inoculation

To evaluate the disease resistance of different rice lines, punch inoculation was carried out using the *Magnaporthe oryzae* Guy11 strain (was kindly provided by Professor Kabin Xie) as described previously (19), with minor modifications. Conidia of *M. oryzae* were harvested from 10-day-old cultures grown on Oat-tomato agar medium (OTA, 30 g oatmeal, 200 mL tomato juice, 20 g agar, 1 L distilled water) under continuous light at 28 °C. The spores were washed from the plate surface with sterile distilled water. The spore concentration was determined with hemocytometers and adjusted to 5 × 10^6^ conidia/mL using sterile distilled water supplemented with 0.02% Tween-20 to promote even suspension.

For inoculation, fully expanded leaves from 4- to 5-week-old rice plants (TN1, 35S::GFP, 35S::BSCL1_wt, and 35S::BSCL1_mut) were excised and placed on moist filter paper inside plastic boxes (30 × 20 × 10 cm). Two small wounds were made on each leaf using a sterile needle, and 5 μL of the spore suspension was carefully pipetted onto each wound site. Each plastic box contained leaves from a single rice line to avoid cross-contamination. After inoculation, the boxes were sealed with plastic bag to maintain high humidity and incubated in the dark at 28 °C for 24 h. Subsequently, the boxes were transferred to growth chambers maintained at 28 °C under a 14 h light / 10 h dark photoperiod. Disease progression was monitored daily. After 6 days, the leaves were photographed using a digital camera, and lesion development was recorded (20, 21). In total, 40 leaves per rice line were inoculated, serving as independent biological replicates for statistical analysis.

##### Gene coexpression analysis and GO enrichment analysis

To investigate the gene expression responses of different rice lines (TN1, 35S::GFP, 35S::BSCL1_wt, and 35S::BSCL1_mut) to BPH feeding, thirty fifth-instar BPH nymphs were confined on the stem of each rice plant at the tillering stage (∼30 days old) using a cylindrical plastic cage. After 24 h of feeding, the outer layer of the feeding site (FS) was carefully excised, immediately snap-frozen in liquid nitrogen, and stored at –80 °C until RNA extraction. Three biological replicates were performed for each rice line, and non-infested plants were included as controls.

RNA sequencing was performed by OE Biotech Co., Ltd. (Shanghai, China). RNA-seq analysis was discribed in the secton "RNA-seq analysis", which included RMTA for reads alignement, HTseq for reads count, and edgeR for identifying DEGs with |log_2_(fold change)| ≥ 1 and FDR-adjusted *p* value < 0.05. Co-expression analysis of DEGs was performed with the Mfuzz package in R (v2.6.1) (22) with default settings, and Gene Ontology (GO) enrichment analysis was carried out using the clusterProfiler package in R (v3.18.1) (23).

##### Localization of *BSCL1* in planta

For visualization of *BSCL1* in plant cells, an RNA-triggered fluorescence (RTF) reporter system was employed (24). The RTF system consists of two modules: (i) an RNA switch containing a probe sequence complementary to the target RNA, and (ii) a GFP expression cassette that is regulated by the RNA switch. In the absence of the target RNA, the RNA switch remains in an inactive conformation, leading to recruitment of the 26S proteasome and degradation of GFP, thereby preventing fluorescence. In contrast, binding of the target RNA to the probe region induces a conformational switch that prevents GFP degradation, allowing GFP accumulation and fluorescence detection.

The RNA switch design method was adapted from Bai et al. (24). Probes were designed using the Stellaris Probe Designer (Biosearch Technologies, https://www.biosearchtech.com/support/tools/design-software/stellaris-probe-designer) with default settings, selecting sequences complementary to *BSCL1* of approximately 21 nt in length and with annealing temperatures below 65 °C. RNA switch secondary structures were predicted using mFold (www.unafold.org/mfold/applications/rna-folding-form.php (25)) with default parameters. Switches adopting a dumbbell-shaped structure, particularly those with shorter stems and larger bottom loops, were critical for functionality and guided the selection of optimal designs (24). The fulll lenght of RNA switch sequeqnce including probe for *BSCL1* was listed in Table S6.

The probe sequence specific for *BSCL1* was cloned into the pcambia1300-AtU6::RNA-Switch–AtUbq10::RTF vector at the designated restriction sites, generating the RNA-switch-*BSCL1*–RTF construct (Fig. S4*E*). The RTF system comprises full-length GFP fused to a single-chain antibody (scFv) carrying an RRRG degron, together with 24 × GCN4 fused to Rev at its N-terminus. In this system, the scFv-GFP-RRRG fusion functions as the reporter, producing fluorescence only when stabilized, while the RRRG degron ensures that unbound GFP is rapidly degraded, minimizing background signal. The 24 × GCN4-Revd2 component binds to the RNA switch containing RRE sequences, recruiting multiple GFP-scFv molecules and thereby amplifying the fluorescence signal at the RNA site. Primers used for cloning are listed in Table S6. The construct was sequence-verified by Sanger sequencing and introduced into *A. tumefaciens* GV3101 (pSoup) (Weidi, AC1002, Shanghai, China).

The RNA-switch-*BCSL1*–RTF construct was co-infiltrated with PC1300S-35S::BSCL1_wt into fully expanded leaves of 4-week-old *N. benthamiana* plants using the leaf infiltration method. *A. tumefaciens* cultures were grown overnight in LB medium containing appropriate antibiotics, harvested by centrifugation, and the bacterial pellet was washed once with infiltration buffer [10 mM MgCl_2_, 10 mM MES pH 5.6 (BioFroxx, 1086GR500, Germany), 100 µM acetosyringone (BioFroxx, 2279GR001, Germany)]. The pellet was then resuspended in infiltration buffer and adjusted to an OD_600_ of 0.6. Equal volumes of GV3101 strains carrying the respective constructs were mixed and incubate in the dark for 2 h prior to infiltration. Infiltration was performed on the abaxial side of leaves using a 1-mL needleless syringe. Plants infiltrated with only the RTF construct or PC1300S-35S::BSCL1_wt served as negative controls, while those infiltrated with two constructs for GFP expression. PC1300S-35S::GFP alone was used as a positive control.

Three days post-infiltration (dpi), infiltrated leaves were harvested, mounted in distilled water on glass slides, and imaged using a Leica SP8 confocal laser scanning microscope (Leica Microsystems, Germany). GFP fluorescence was excited at 488 nm and detected between 500–530 nm. Chlorophyll autofluorescence was collected at 650–700 nm to assist with subcellular localization.

##### Yeast three-hybrid (Y3H) assay

Y3H was adopted from the protocol developed by Professor Marvin Wickens’ lab at the University of Wisconsin-Madison (26). The plasmid, p3HR2 for expressing *BSCL1*, pIIIA/IRE-MS2 and pAD-IRP severed as positive controls, the yeast strain YBZ1, were kindly provided by Professor Marvin Wickens.

The RNA expression plasmid was generated by inserting the 708-nt *BSCL1* sequence into p3HR2 between XhoⅠ and XmaⅠ restriction sites. The recombinant plasmid p3HR2-*BSCL1* was verified by restriction digestion and sequencing after amplification in *E. coli* (primers were listed in Table S6). The *Saccharomyces cerevisiae* strain YBZ1 was used for yeast three-hybrid assays. Competent cells were prepared as follows: a single colony was inoculated from a YPDA (Coolaber, PM2011, Beijing, China) plate into 5 mL YPDA broth medium and grown overnight at 30 °C with shaking (250 rpm). Cells were diluted to an OD600 of 1.2–1.5, pelleted at 1,000 g for 5 min, and sequentially washed with sterile water and TE/LiAc buffer [1 mL 10 × TE, 1 mL 10 × LiAc (Sigma, L4158, USA), 8 mL sterile water]. The final cell pellet was resuspended in 1× TE/LiAc to obtain competent cells.

Transformation was performed by mixing competent cells with 40% PEG4000/TE/LiAc [1 mL 10 × TE, 1 mL 10 × LiAc (Sigma, L4158, USA), 8 mL 50% PEG4000 (Sigma, 95904, USA)], denatured salmon sperm DNA (Thermo Fisher Scientific, 15632011, USA), and p3HR2-*BSCL1*, followed by incubation at 30 °C for 30 min and heat shock at 42 °C for 10 min. After recovery, cells were plated onto synthetic defined (SD) agar medium (Coolaber, PM2040, Beijing, China) lacking uracil (Coolaber, PM2270, Beijing, China) (SD-U) and incubated at 30 °C for 2–3 days. Single colonies were verified by plasmid rescue, restriction digestion, and sequencing. Competent cells containing the p3HR2-*BSCL1* were used for library transformation.

To optimize 3-amino-1,2,4-triazole (3-AT) (Sigma, A8056, USA) concentration and test for autoactivation, yeast strains harboring RNA plasmids were co-transformed with empty pGAD-T7 vector. Transformants were spotted or plated onto SD agar medium (Coolaber, PM2040, Beijing, China) lacking uracil, leucine, and histidine (Coolaber, PM2170, Beijing, China) (SD-U-L-H), supplemented with increasing concentrations of 3-AT (0–100 mM). Growth was monitored after 3–5 days at 30 °C, and the minimal concentration of 3-AT that suppressed background growth was selected for subsequent screening.

Competent cells containing the p3HR2-*BSCL1* were prepared as described above. For library transformation, cells were mixed with 40% PEG4000/TE/LiAc (1 mL 10 × TE, 1 mL 10 × LiAc, 8 mL 50% PEG4000), denatured salmon sperm DNA (Thermo Fisher Scientific, 15632011, USA), and 20 µg of cDNA library, kindly provided by Professor Yongjun Lin at Huazhong Agricultural University. After incubation at 30 °C for 30 min with shaking, DMSO was added, and cells were heat shocked at 42 °C for 15 min. Transformants were plated on SD-U-L-H plates supplemented with the optimized concentration of 3-AT. After 5–7 days at 30 °C, colonies growing on selective plates were isolated as HIS^+^ candidates.

##### LacZ reporter assay and identificaiton of putative interactors

The methods for LacZ screening and interaction validation were adapted from Bernstein et al. (27). Briefly, HIS^+^ colonies were replated onto SD agar medium (Coolaber, PM2040, Beijing, China) lacking uracil and leucine (Coolaber, PM2290, Beijing, China) (SD-U-L), overlaid with nitrocellulose membranes. After incubation at 30 °C for 2–3 days, the membranes were briefly frozen in liquid nitrogen and then incubated at 37 °C in buffer (Z-buffer supplemented with 65 µL β-mercaptoethanol and 200 µL of 25 mg/mL X-β-gal per 10 mL). Colonies that developed a blue color were scored as LacZ-positive.

Plasmids were extracted from LacZ-positive yeast colonies, and the inserted sequences were amplified by PCR, verified by Sanger sequencing, and identified through BLAST searches against the NCBI database. To validate *BSCL1*–protein interactions, full-length cDNAs of candidate genes were cloned into pGAD-T7 and co-transformed into YBZ1 cells together with either p3HR2-BSCL1 or the empty p3HR2 vector (negative control). The pIIIA/IRE-MS2 and pAD-IRP plasmids served as positive controls. Transformants were plated onto SD-U-L-H medium supplemented with the optimized concentration of 3-AT. Growth observed with the *BSCL1* plasmid but absent with the empty vector was taken as evidence of a specific RNA–protein interaction. Colonies that turned blue on SD-U-L medium in the X-β-gal reporter assay were considered to represent positive interactions.

##### Co-localization of *BSCL1* with candidate target proteins

The full-length CDSs of HIRIP3 and H3.3 were amplified by RT-PCR from rice cDNA and cloned into the pB7RWG between SpeⅠ and XhoⅠ restriction sites to generate C-terminal fusions with mCherry. Constructs were verified by Sanger sequencing and introduced into *A. tumefaciens* GV3101 (pSoup) (Weidi, AC1002, Shanghai, China) as described above.

*Agrobacterium* strains carrying pB7RWG-HIRIP3-mCherry or pB7RWG-H3.3-mCherry were co-infiltrated with PC1300S-35S::BSCL1_wt and the RNA-switch-*BSCL1*–RTF construct into *N. benthamiana* leaves, following the same infiltration protocol as above. Each protein was tested in independent infiltration assays.

At 3 dpi, infiltrated leaves were examined by confocal microscopy Leica SP8 (Leica Microsystems, Germany). GFP signals (*BSCL1* detection via RNA-switch-*BSCL1*–RTF system) were excited at 488 nm (emission 500–530 nm), while mCherry-tagged HIRIP3 and H3.3 were excited at 561 nm (emission 580–610 nm). Chlorophyll autofluorescence was collected at 650–700 nm to assist with subcellular localization. Images were collected in sequential scanning mode to prevent bleed-through between channels. Representative images were captured and processed using identical microscope settings for all samples.

##### dsRNA synthesis, microinjection, and gene expression analysis in BPH

Double-stranded RNA (dsRNA) targeting *BSCL1* was synthesized by PCR amplification of two gene fragments (*dsBSCL1*-F1/*dsBSCL1*-R1: 315 bp, *dsBSCL1*-F2/*dsBSCL1*-R2: 260 bp) with primers containing T7 RNA polymerase promoter sequences at both 5’ ends (Table S6). The PCR products were used as templates for in vitro transcription with the T7 High Yield RNA Transcription Kit (Vazyme, TR101-01, Nanjing, China) following the manufacturer’s instructions. Synthesized dsRNAs were diluted to the appropriate concentration for microinjection.

Third- to fourth-instar BPH nymphs were anesthetized with carbon dioxide for 20 s and injected with 200 ng dsRNA into the mesothorax using a Nanoliter 2010 microinjector (World Precision Instruments, USA). BPH injected with *dsGFP* served as the negative control. Each treatment was performed in three biological replicates. For RNAi efficiency assessment, ten nymphs per replicate were collected at 3 days post-injection. The remaining injected nymphs were maintained on three-leaf stage rice seedlings for subsequent experiments.

Total RNA was isolated from microinjected nymphs or insects at different developmental stages using TRIzol reagent (Invitrogen, 15596018CN, USA). First-strand cDNA synthesis was performed with the PrimeScript RT Reagent Kit with gDNA Eraser (Takara, RR047A, Japan) following the manufacturer’s instructions. qRT-PCR was conducted as described above using gene-specific primers, with *N. lugens Actin* (*NlActin*) serving as the endogenous control (primers were listed in Table S6). Relative expression levels were calculated using the 2^^-ΔΔCt^ method.

##### BPH survival and fecundity assays

To evaluate the effects of dsRNA-mediated gene silencing on BPH survival, approximately 100 third- to fourth-instar nymphs treated with gene-specific or control dsRNAs were transferred into 500-mL glass beakers containing fresh three-leaf stage TN1 seedlings (28). Each beaker was treated as one biological replicate, with three replicates per treatment. Rice seedlings were replaced every 4 days to ensure a continuous food supply and minimize confounding effects from plant senescence. Beakers were maintained in a controlled growth chamber at 27 ± 1 °C, 70 ± 5 % relative humidity, and a 14:10 h light:dark photoperiod. The number of surviving nymphs was recorded every 24 h, and mortality was calculated relative to the initial number of insects. Monitoring continued until most individuals reached adulthood or exhibited mortality consistent with RNAi effects.

Fecundity was assessed according to the methods described by Wen et al. (29) and Liu et al. (30), with minor modifications. For BPH fecundity assays following microinjection, newly emerged dsRNA-treated adults (≤ 24 h old) were paired in glass tube (3 cm diameter, 25 cm high) supplied with five three-leaf stage TN1 rice seedlings for feeding and oviposition. Adults were transferred to a new glass tube with fresh rice seedlings every 4 days until all of the adults were dead. For fecundity assays on different rice lines, each rice plant was enclosed with a perforated plastic lid to allow plant growth and covered with an inverted transparent plastic cup (7.5 cm diameter) to confine insects. Five newly hatched nymphs (≤ 24 h old) were placed into each plastic cup containing a single rice plant. Upon adult emergence, one male–female pair was retained per cup for oviposition and removed after 10 days. Newly hatched nymphs were counted and removed daily until no further hatching occurred. A minimum of 15 biological replicates were performed for each dsRNA treatment or rice line to ensure robust statistical analysis.

##### Honeydew secretion assay

Honeydew secretion was measured following the method described by Guo et al. (31). As described in fecundity assays, each rice plant was covered with a perforated plastic lid that allowed the plant to grow through. A 9-cm diameter filter paper was placed on the lid and covered with an inverted transparent plastic cup (7.5 cm diameter) to confine the insects.

For assays after microinjection, two dsRNA-treated 5th instar nymphs were introduced into each cup and allowed to feed for 48 h, after which filter papers were collected. At least 15 biological replicates were performed per treatment. For assays comparing BPH feeding on different rice lines, five 5th instar nymphs were confined per cup and allowed to feed for 48 h, with at least 20 replicates per rice line.

Collected filter papers were soaked in 0.1% (w/v) ninhydrin in acetone, dried in an oven at 60 °C for 30 min, and honeydew spots visualized as violet or purple stains due to their amino acid content. Images were captured using a digital camera, and honeydew spot areas were quantified using ImageJ software.

##### Dual-luciferase reporter assay in *N. benthamiana*

Dual-luciferase reporter assays were performed as previously described (32). The 1989-bp promoter region of bHLH6 was amplified from rice genomic DNA by PCR and cloned upstream of the firefly luciferase (LUC) coding sequence in the pGreenⅡ 0800-LUC vector, generating the Pro_bHLH6_::LUC reporter construct. In this vector, renilla luciferase (REN) driven by the CaMV 35S promoter served as an internal control for normalization of LUC activity.

For effector constructs, the full-length sequence of *BSCL1_wt*, the full-length coding sequences (CDSs) of *HIRIP3* and *H3.3*, as well as a truncated *GFP* sequence (negative control for *BSCL1*_wt), were amplified by PCR and cloned separately into pGreenⅡ 62-SK under the control of the CaMV 35S promoter. Primers used for all constructs are listed in Table S6. All plasmids were verified by Sanger sequencing using specific primers and introduced into *A. tumefaciens* GV3101 (pSoup) (Weidi, AC1002, Shanghai, China).

*Agrobacterium* strains carrying the reporter and effector constructs were cultured overnight in LB medium containing appropriate antibiotics at 28 °C with shaking. Cells were harvested by centrifugation, washed, and resuspended in infiltration buffer [10 mM MgCl₂, 10 mM MES pH 5.6 (BioFroxx, 1086GR500, Germany), 100 µM acetosyringone (BioFroxx, 2279GR001, Germany)] to an OD_600_ of 0.6. For co-infiltration, equal volumes of *Agrobacterium* cultures carrying reporter and effector constructs were mixed. Fully expanded leaves of 4-week-old *N. benthamiana* plants were infiltrated on the abaxial side using a 1-mL needleless syringe. Each construct combination was infiltrated into at least three independent leaves from different plants as biological replicates.

Infiltrated plants were maintained under standard growth conditions (22–25 °C, 14 h light/10 h dark) for 3 days to allow transient expression. Firefly (LUC) and renilla (REN) luciferase signals were measured using the Dual-Luciferase Reporter Assay Kit (Vazyme, DL101-01, Nanjing, China) according to the manufacturer’s instructions. Leaf discs (approximately 1.0 cm diameter) were collected from infiltrated zones, homogenized in passive lysis buffer, and luminescence was measured with a multimode reader Spark (TECAN, Zurich, Switzerland). Relative LUC activity was calculated as the ratio of LUC:REN signals for each sample.

For live imaging, D-Luciferin potassium salt (1 mM, Yeasen, 40902ES01, Shanghai, China) was evenly applied to the abaxial surface of infiltrated leaves. Luminescence was visualized using the NightSHADE L985 live imaging system (Berthold, Germany), and images were captured under standardized exposure conditions.

##### Data analysis

Statistical analysis was performed with SPSS 20.0 (IBM, Armonk City, NY, USA). Paired Student’s *t* test or one-way ANOVA test followed by Tukey’s multiple comparisons test was used to analyze the results of survival rate, honeydew measurement, fecundity measurement, and qRT-PCR. Correlation analysis and Fisher’s Exact test were performed in R. *P* < 0.05 was considered as statistically significant. All results are presented as means ± SE.

---

**Fig. S1.**
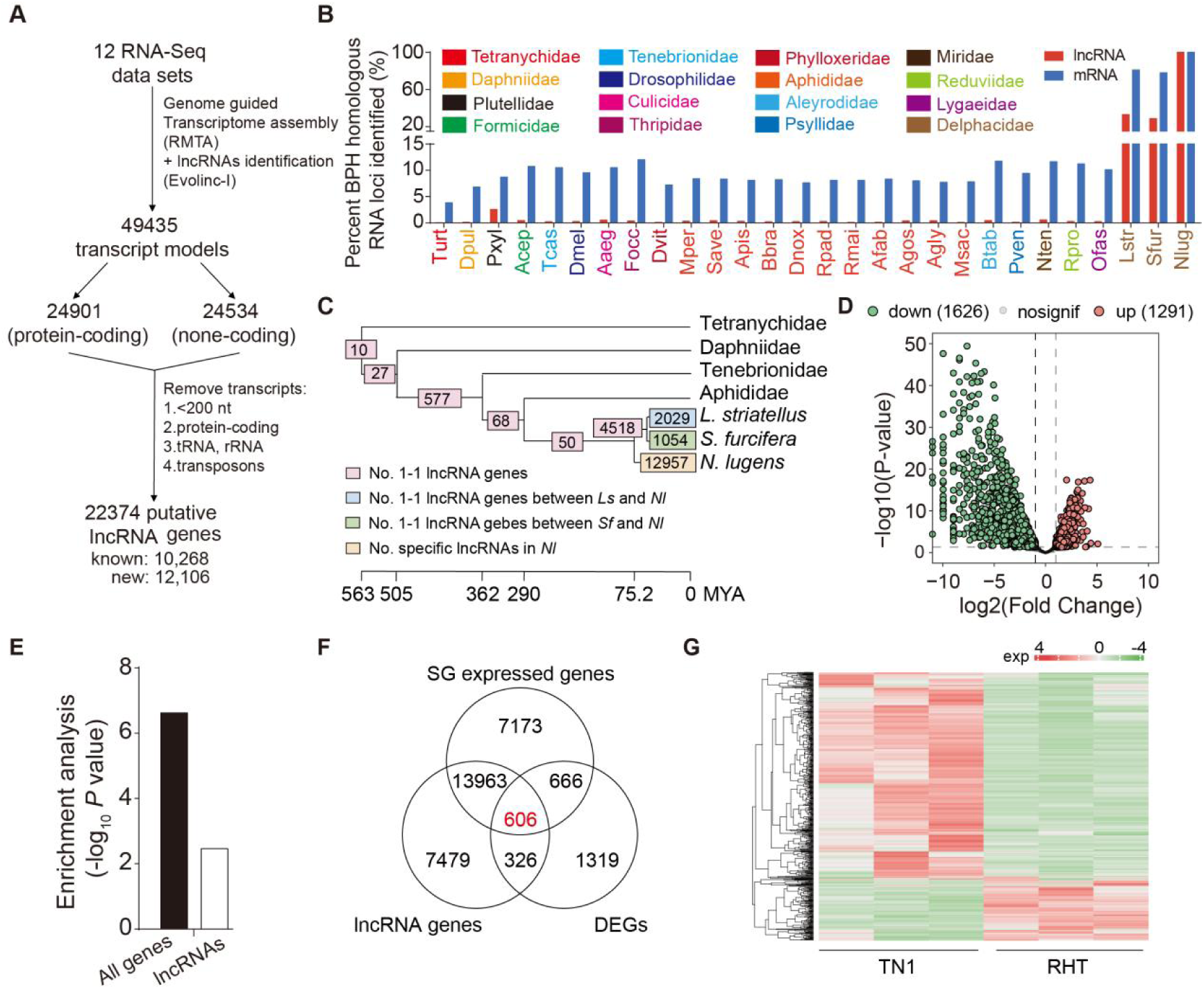
Identification of lncRNA genes BPH and anlaysis their resposne to resistant rice. (A) Workflow for the identification of lncRNAs in BPH (For details, please refer to the Materials and Methods section). (B) Protein-coding genes (mRNAs) exhibit higher conservation than lncRNAs across 28 species. Detailed species information and data sources are listed in Table S5. Red and blue bars indicate the percentages of BPH-homologous lncRNA and mRNA loci identified in each species, respectively. Percentage of BPH-homologous loci in each species was calculated as the number of homologs divided by total BPH transcripts. (C) Phylogenetic distribution of 1:1 orthologous and species-specific lncRNAs in BPH. Numbers in the pink boxes represent lncRNAs homologous to BPH in other species outside Delphacidae across different evolutionary periods; numbers in the blue and green boxes indicate lncRNAs homologous to BPH in the *L. striatellus* and *S. furcifera*, respectively; number in the yellow box represent BPH-specific lncRNAs. (D) Volcano plot of DEGs in BPH feeding on resistant rice RHT versus susceptible rice TN1. (E) Salivary gland was enriched for DE genes and DE lncRNAs in BPH. Fisher’s exact test was used to assess the enrichment of DE genes and DE lncRNAs in the salivary gland. For each category, the number expressed in the salivary gland was compared to the total number of DE genes or DE lncRNAs. Enrichment with *P* < 0.05 was considered statistically significant. Black bars represent all DEGs; white bars, candidate lncRNAs. (F) Venn diagram showing the overlap of lncRNAs identified in BPH, DEGs in response to RHT, and genes expressed in BPH salivary gland. (G) Heatmap of transcript abundance (TPM values) corresponding to the genes in (F).

**Fig. S2.**
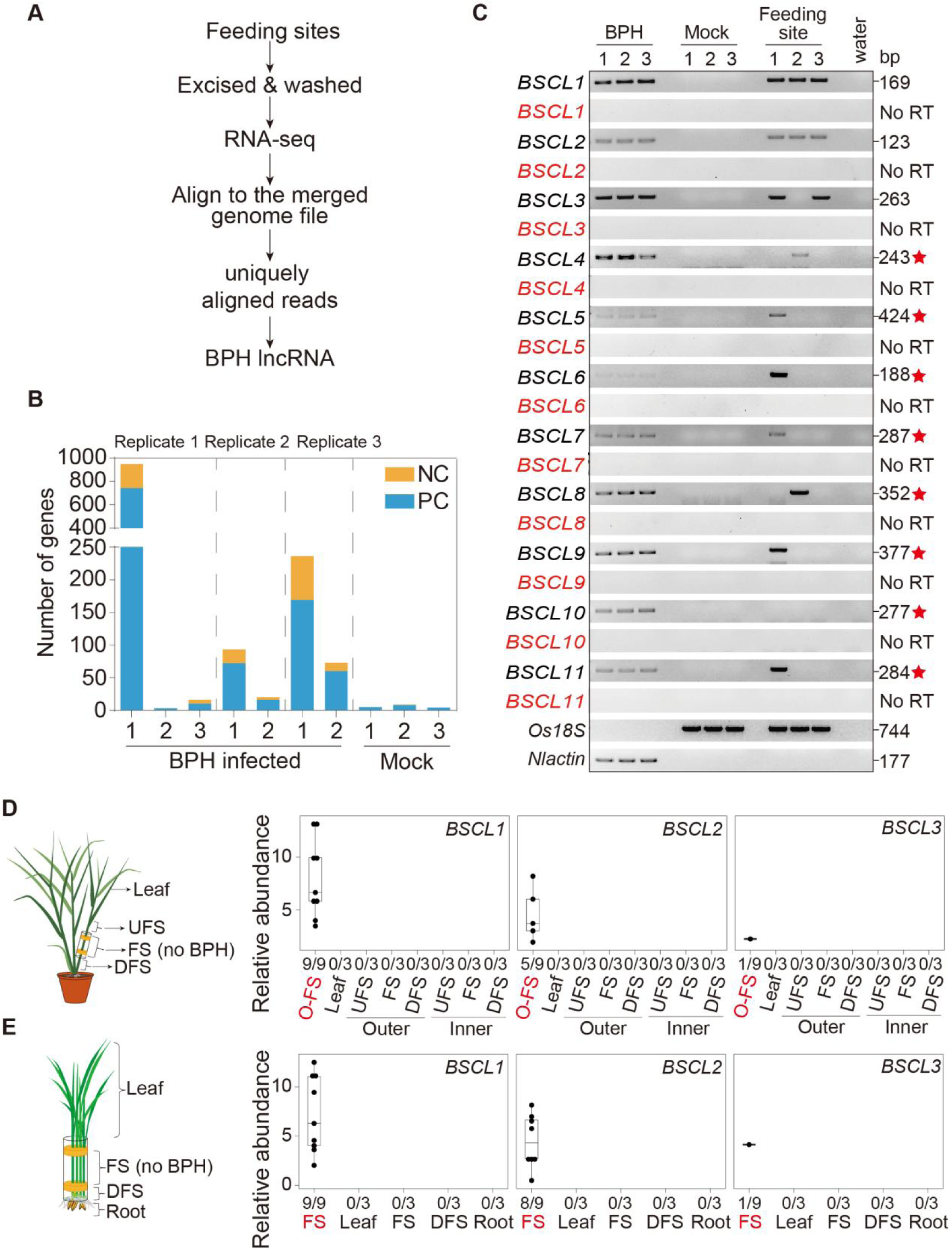
Identification of BPH-delivered RNAs in rice plants. (A) Workflow for the identification of BPH-delivered RNA transcripts in FS (For details, please refer to the Materials and Methods section). (B) Numbers of BPH-delivered RNA transcripts identified across different samples. NC, none coding transcripts; PC, protein coding transcripts. (C) *BSCL*s were translocated into the FS of rice plants. No RT indicates PCR performed using RNA as the template without reverse transcription. Red stars denote genes subjected to a second round of PCR using PCR products as templates. (D) *BSCL1*, *BSCL2*, and *BSCL3* were undetectable in leaf blade, FS, UFS, and DFS regions in uninfested tillering stage rice plants. A schematic of leaf blade, FS, UFS, and DFS is shown on the left. Box plots indicate the relative abundance of *BSCL1*, *BSCL2*, and *BSCL3* at different sampling sites, determined by qRT-PCR. (E) *BSCL1*, *BSCL2*, and *BSCL3* were undetectable in leaf, FS, DFS, and root regions in uninfested three-leaf stage rice plants. A schematic of leaf blade, FS, DFS and root is shown on the left. Box plots indicate the relative abundance of *BSCL1*, *BSCL2*, and *BSCL3* at different sampling sites, determined by qRT-PCR. Numbers under the boxpot show the detection ratio of *BSCL1, BSCL2* and *BSCL3* at each site. The sampling sites marked in red are from samples that were fed by BPH, consistent with those in Fig. 1.

**Fig. S3.**
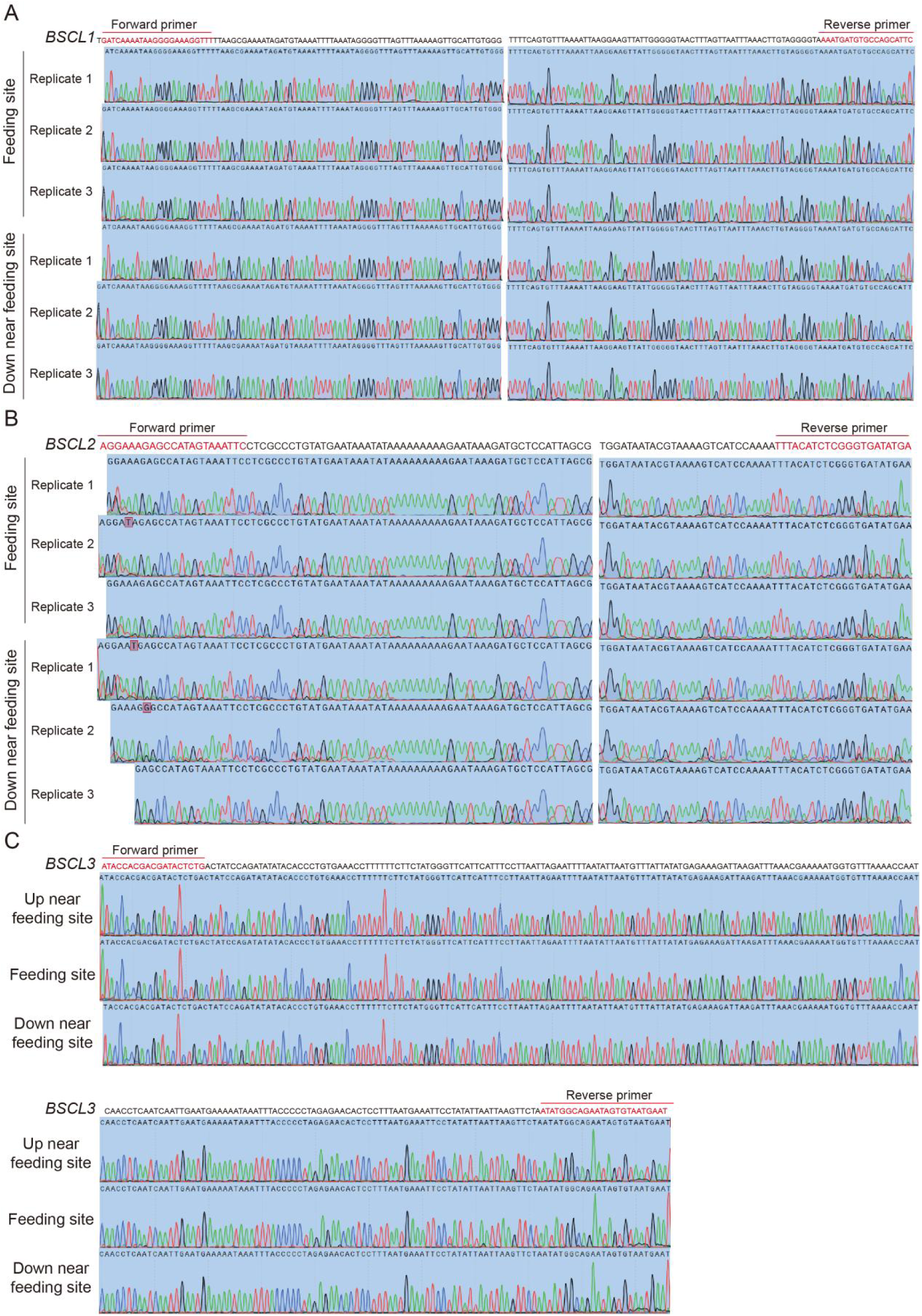
Sequences of BPH *BSCLs* amplified from rice plants via RT-PCR. The cDNA sequence of *BSCL1* (A), *BSCL2* (B), and *BSCL3* (C) were amplified from FS, DFS, and UFS of rice plants with corresponding forward and reverse primers (Primer sequences are listed in Table S6). The sequence of PCR products were identical to the corresponding regions of their respective *BSCL*s.

**Fig. S4.**
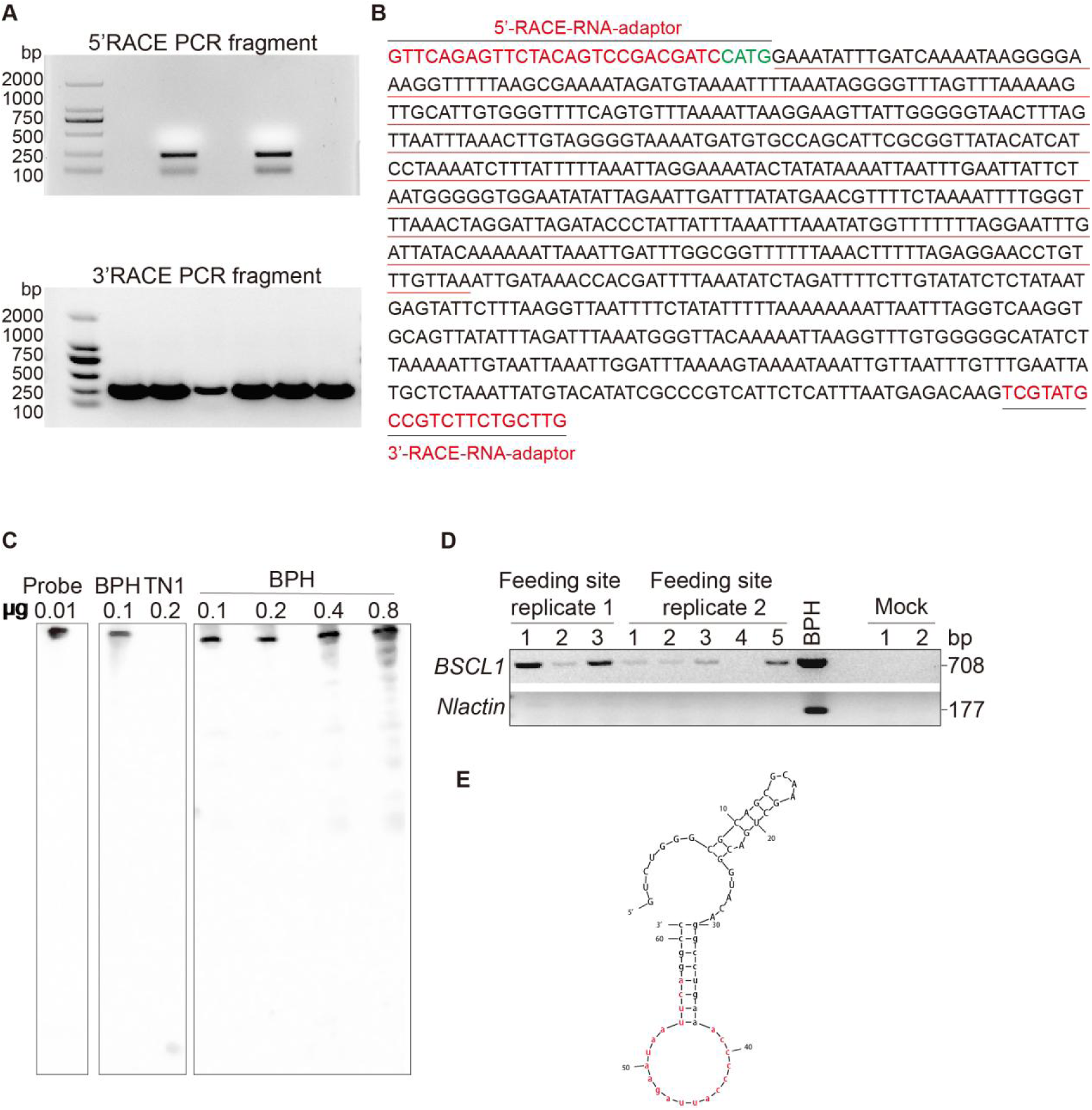
*BSCL1* is secreted into rice in the form of full-length transcripts. (A) Results of a 5’ and 3’ RACE experiment. The PCR products were separated on a 2% agarose gel, and the target bands were excised, purified, and sequenced. The resulting sequences were then assembled with the predicted *BSCL1* sequence. (B) Sequence of the *BSCL1* full-length cDNA. The sequences marked in red at both ends represent 5’ and 3’ RNA adapter sequences, respectively. The sequence underlined in red corresponds to the annotated *BSCL1* sequence, and the sequence downstream of its 3’ end was obtained from the 3’ RACE experiment. (C) Northern blot hybridizations with full length *BSCL1* probe showing no obvious short fragment in BPH RNAs. (D) Full-length transcript of *BSCL1* was detected in FS via RT-PCR. Numbers above the gel image were the number of samples used in each replicate. (E) The secondary structure of RNA-switch was designed for targeting of *BSCL1*. The sequences marked in red represent the probe specifically binding to *BSCL1* (Predicted free energy: dG = -21.70 kcal/mol), and the design principles and screening criteria of the RNA-switch are described in the Materials and Methods section.

**Fig. S5.**
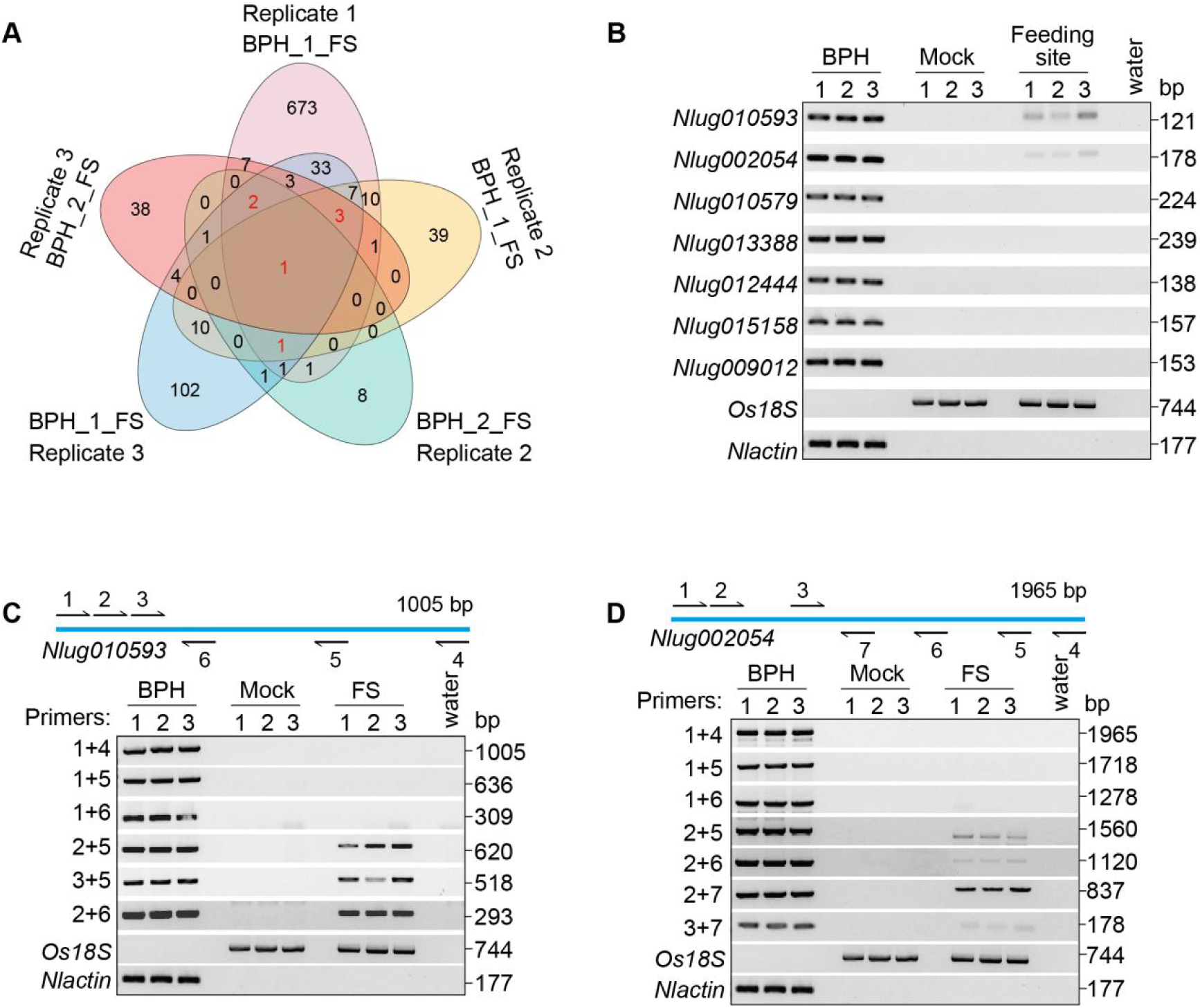
BPH mRNAs are transcloated into rice plants. (A) Venn diagram showing the numbers of BPH mRNAs identified in FS samples. (B) BPH mRNA (*Nlug010593* and *Nlug002054*) transcripts were translocated into the FS of rice plants. (C, D) Identification of *Nlug010593* (C) and *Nlug002054* (D) transcripts sizes in BPH and FS. Schematics above the gel images indicate the locations of primers used to detect *Nlug010593* (primers 1–6) and *Nlug002054* (primers 1–7) transcripts in BPH and FS by RT-PCR, shown as black arrows. The full-lenghth of *Nlug010593* (C) and *Nlug002054* (D) transcripts were detected in BPH and not in FS, where the shorter fragements were found in FS.

**Fig. S6.**
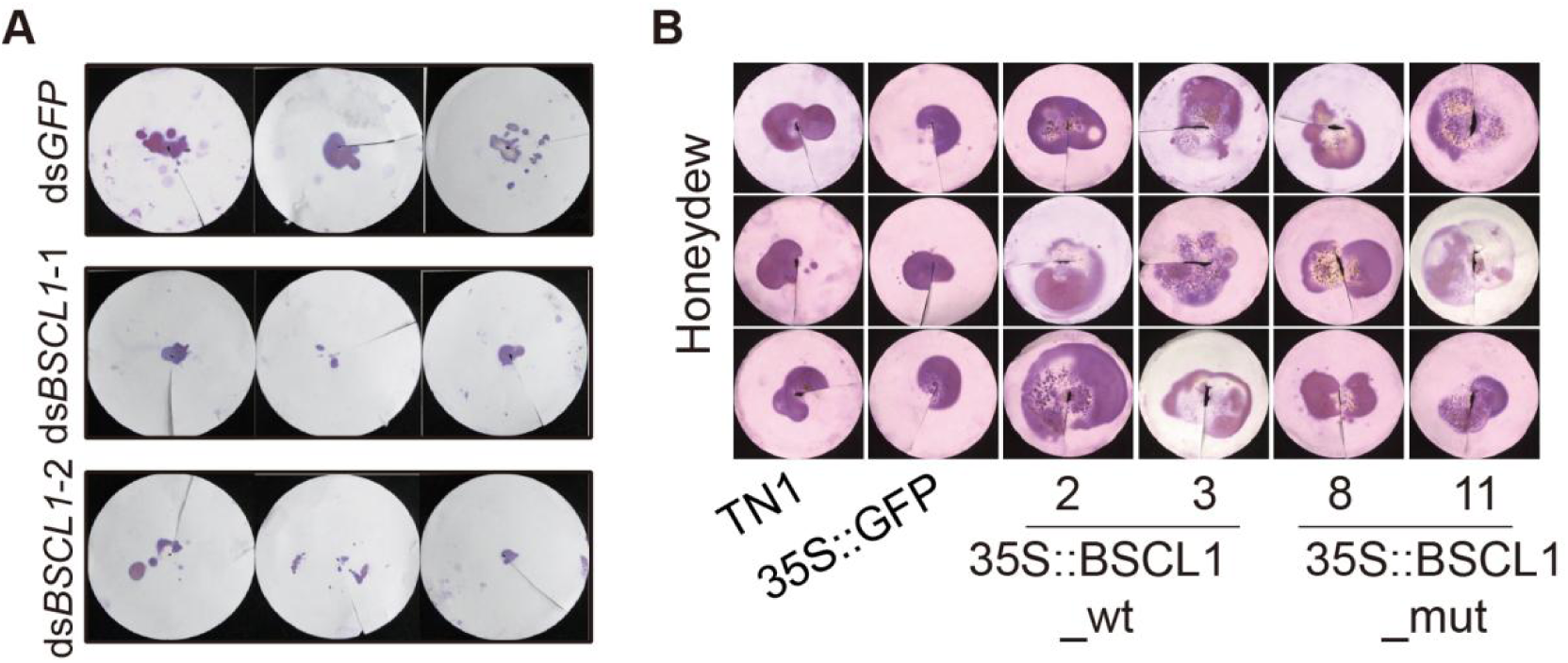
Detection of brown planthopper honeydew secretion after *BSCL1* interference (A) and after feeding on different rice lines (B).

**Fig. S7.**
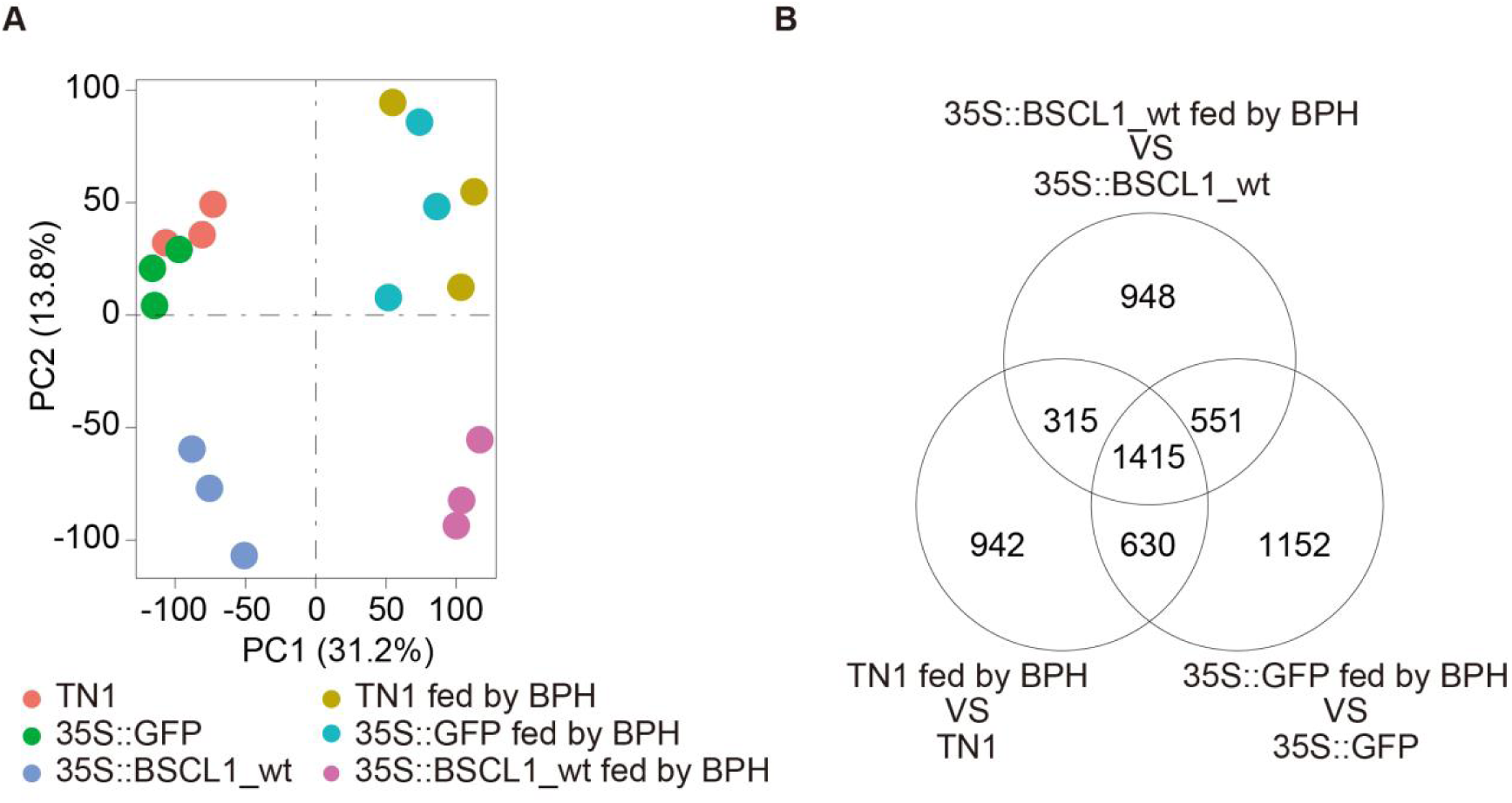
*BSCL1*-overexpressing rice shows a distinct gene expression pattern from TN1 and GFP-overexpressing rice. (A) PCA of RNA-seq data reveals distinct clustering of rice lines with or without 24 h BPH feeding. Three biological replicates were included. PC1 and PC2 represent the first and second principal components. (B) Venn diagram showing the numbers of DEGs among different rice lines with or without 24 h BPH feeding.

**Fig. S8.**
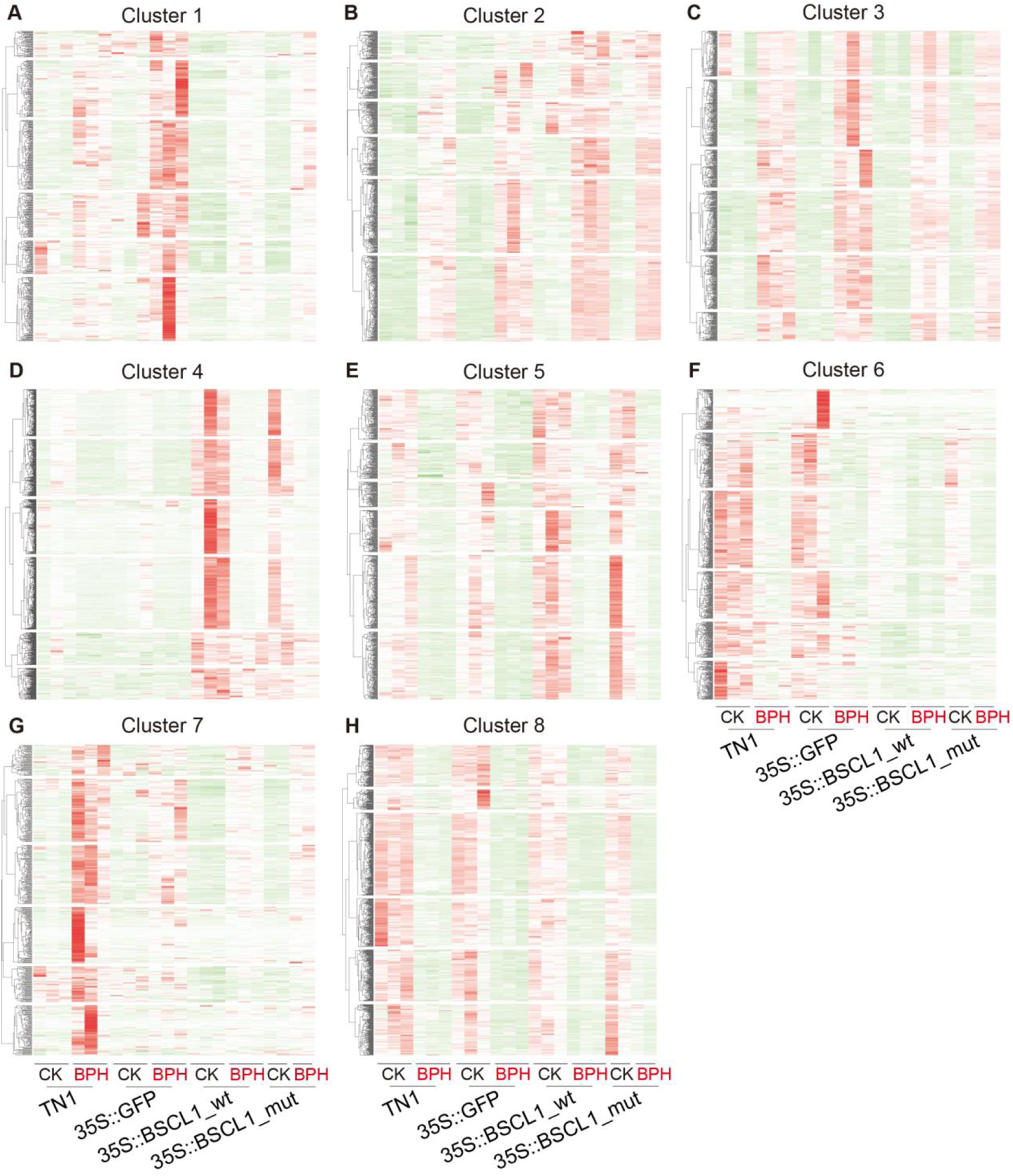
Heatmap of TPM values for DE genes in the clusters. Coexpression patterns DE genes in the clusters were shown in Fig. 5A. CK, uninfested plants; BPH, plants infested by BPH. All of the DE genes from different clusters are listed in Dataset S5.

**Fig. S9.**
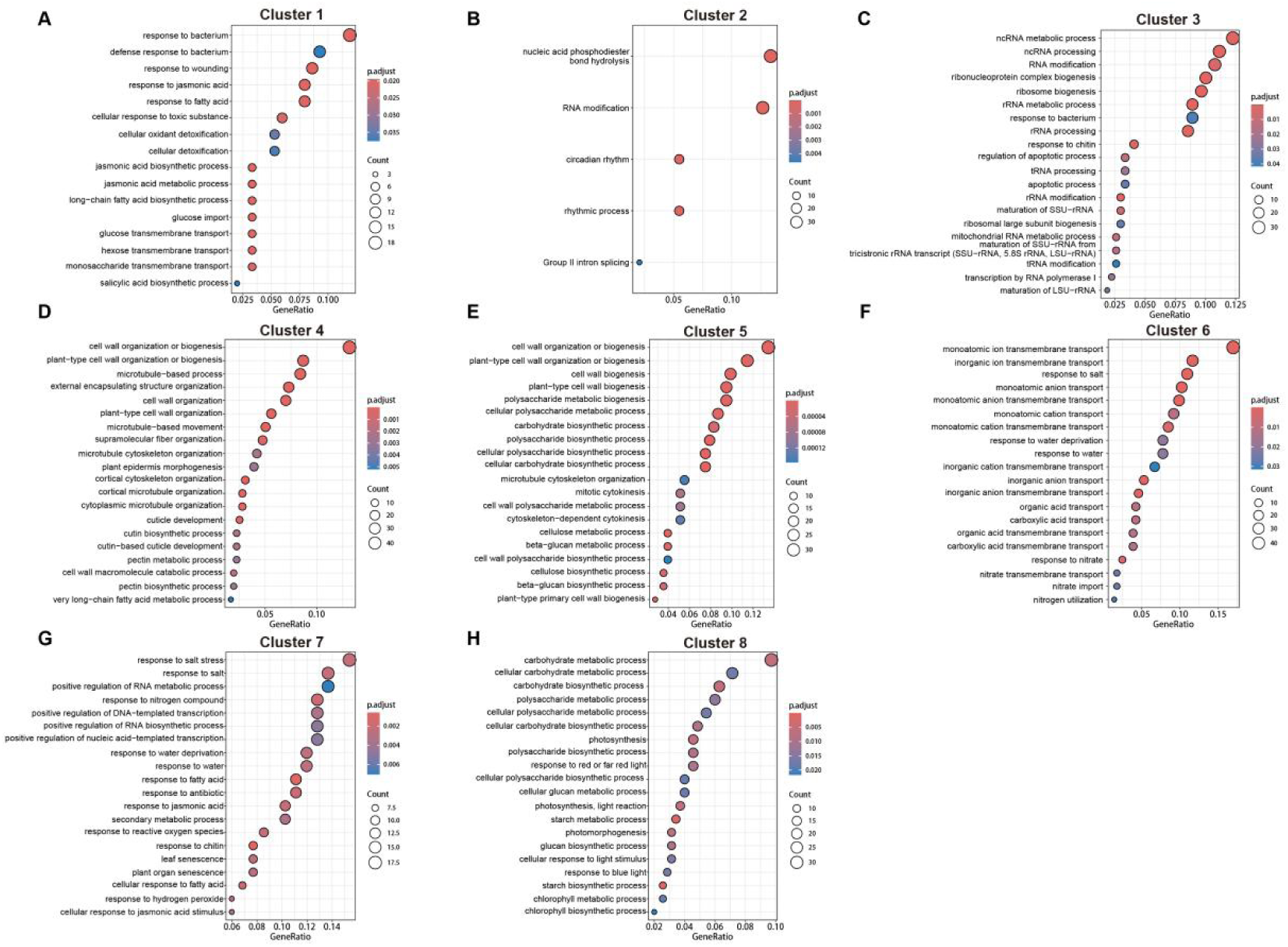
GO enrichment analysis of the eight DE gene clusters. Coexpression patterns DE genes in the clusters were shown in Fig. 5A. Each cluster was analyzed separately to identify significantly enriched GO terms. All DE genes from the clusters are listed in Dataset S5.

**Fig. S10.**
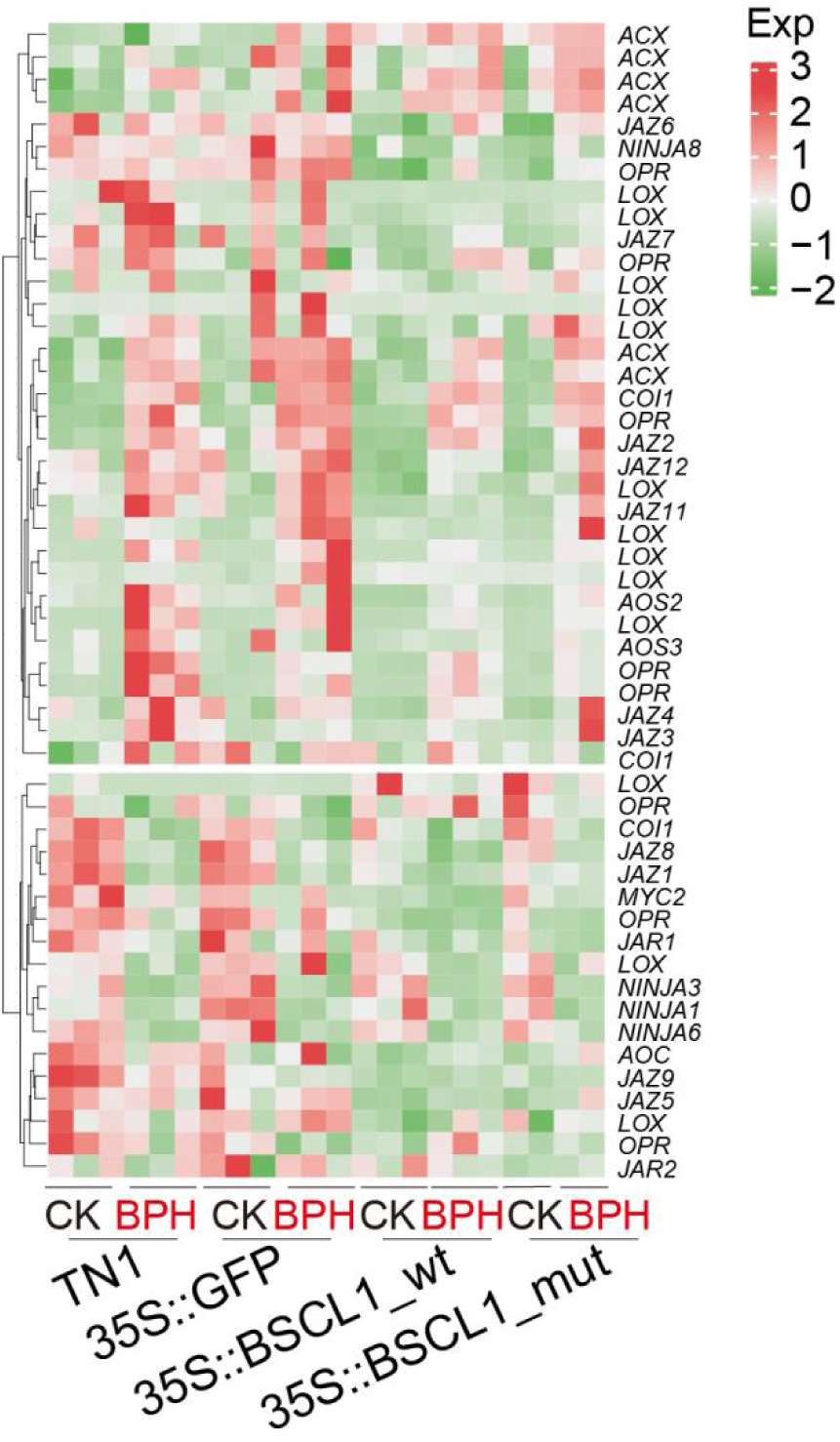
Heatmap of TPM values for genes involved in the JA pathway. CK, uninfested plants; BPH, plants infested by BPH.

**Fig. S11.**
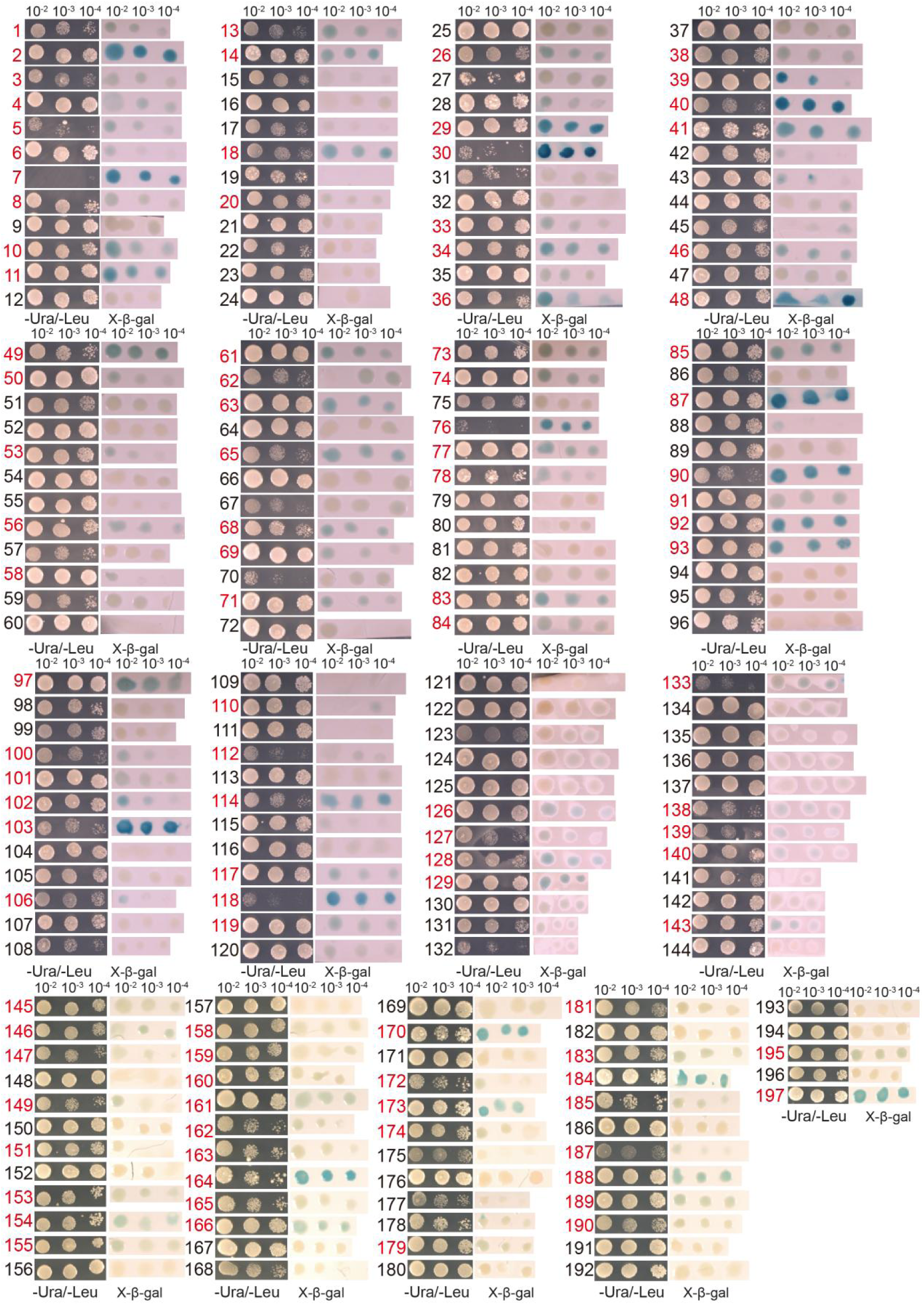
Yeast three-hybrid screening for *BSCL1*-interacting proteins in rice. The obtained monoclonal colonies were diluted 100-, 1000-, and 10,000-fold, respectively, and cultured on SD-U-L medium followed by LacZ reporter assay (For details, please refer to the Materials and Methods section). The red-labeled numbers indicate candidate positive colonies that turned blue in the X-β-gal screening.

## Supplementary tables

**Table S1.**
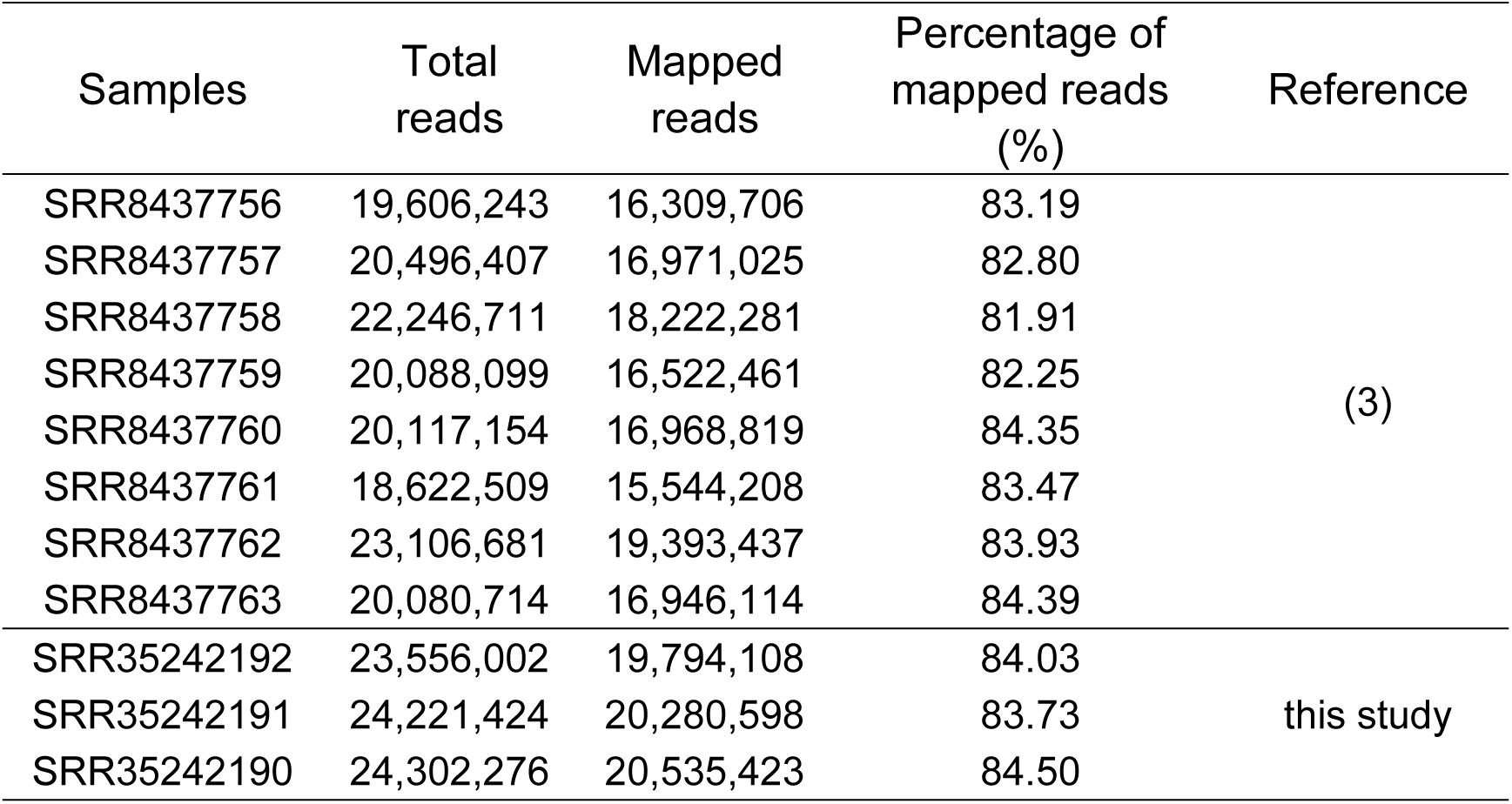
List of RNA-seq data used for BPH lncRNA identification.

| Samples | Total reads | Mapped reads | Percentage of mapped reads (%) | Reference |
| --- | --- | --- | --- | --- |
| SRR8437756 | 19,606,243 | 16,309,706 | 83.19 | (3) |
| SRR8437757 | 20,496,407 | 16,971,025 | 82.80 |  |
| SRR8437758 | 22,246,711 | 18,222,281 | 81.91 |  |
| SRR8437759 | 20,088,099 | 16,522,461 | 82.25 |  |
| SRR8437760 | 20,117,154 | 16,968,819 | 84.35 |  |
| SRR8437761 | 18,622,509 | 15,544,208 | 83.47 |  |
| SRR8437762 | 23,106,681 | 19,393,437 | 83.93 |  |
| SRR8437763 | 20,080,714 | 16,946,114 | 84.39 |  |
| SRR35242192 | 23,556,002 | 19,794,108 | 84.03 | this study |
| SRR35242191 | 24,221,424 | 20,280,598 | 83.73 |  |
| SRR35242190 | 24,302,276 | 20,535,423 | 84.50 |  |

**Table S2.** BPH-delivered mRNAs detected in feeding sites. The samples from different replicates were consistent with the Dataset S2

| Gene_id | Reads | Describe | Replicate 1 | Replicate 2 |  | Replicate 3 |  |
| --- | --- | --- | --- | --- | --- | --- | --- |
|  |  |  | BPH_1_FS | BPH_1_FS | BPH_2_FS | BPH_1_FS | BPH_2_FS |
| *Nlug010593* | 10 | Elongation factor 1-alpha 2 | √ | √ |  | √ | √ |
| *Nlug002054* | 6 | Heat shock 70 kDa protein cognate 4 | √ | √ | √ | √ |  |
| *Nlug010579* | 4 | Soma ferritin | √ | √ |  | √ | √ |
| *Nlug013388* | 21 | Fatty acid synthase | √ | √ | √ | √ | √ |
| *Nlug012444* | 4 | ATP synthase lipid-binding protein, mitochondrial | √ |  | √ | √ | √ |
| *Nlug015158* | 10 | Eukaryotic translation elongation factor 2 | √ |  | √ | √ | √ |
| *Nlug009012* | 18 | Actin-5C | √ | √ |  | √ | √ |

**Table S3.** List of enriched genes from transcription factors WRKY and bHLH families in clusters 1, 7, 3, and 6.

| Gene_id | Gene_name | Cluster |
| --- | --- | --- |
| *OsTN11t000130* | *WRKY70* | Cluster1 |
| *OsTN11t000133* | *WRKY70* | Cluster1 |
| *OsTN11t000134* | *WRKY27* | Cluster1 |
| *OsTN11t001671* | *WRKY72* | Cluster1 |
| *OsTN12t000123* | *WRKY63* | Cluster1 |
| *OsTN12t000129* | *WRKY72* | Cluster1 |
| *OsTN1t001354* | *WRKY5* | Cluster1 |
| *OsTN1t003729* | *WRKY19* | Cluster1 |
| *OsTN5t000974* | *WRKY70* | Cluster1 |
| *OsTN5t002233* | *WRKY55* | Cluster1 |
| *OsTN5t002805* | *WRKY23* | Cluster1 |
| *OsTN9t001168* | *WRKY76* | Cluster1 |
| *OsTN9t001169* | *WRKY62* | Cluster1 |
| *OsTN1t000645* | *WRKY72* | Cluster7 |
| *OsTN1t000646* | *WRKY50* | Cluster7 |
| *OsTN1t002732* | *WRKY16* | Cluster7 |
| *OsTN1t003730* | *WRKY45* | Cluster7 |
| *OsTN1t003764* | *WRKY24* | Cluster7 |
| *OsTN2t000659* | *WRKY71* | Cluster7 |
| *OsTN4t000802* | *WRKY51* | Cluster7 |
| *OsTN5t001375* | *WRKY45* | Cluster7 |
| *OsTN6t002598* | *WRKY28* | Cluster7 |
| *OsTN8t000571* | *ZBED* | Cluster7 |
| *OsTN8t001365* | *WRKY19* | Cluster7 |
| *OsTN9t000735* | *WRKY74* | Cluster7 |
| *OsTN1g002190* | *bHLH041* | Cluster3 |
| *OsTN10g001009* | *bHLH96* | Cluster6 |
| *OsTN12g002074* | *PIF13* | Cluster6 |
| *OsTN1g002203* | *R-S PROTEIN* | Cluster6 |
| *OsTN1g002983* | *bHLH13* | Cluster6 |
| *OsTN1g004238* | *bHLH128* | Cluster6 |
| *OsTN2g003177* | *bHLH79* | Cluster6 |
| *OsTN2g003535* | *bHLH94* | Cluster6 |
| *OsTN4g000845* | *bHLH6* | Cluster6 |
| *OsTN8g002021* | *bHLH130* | Cluster6 |
| *OsTN9g001166* | *RHL1* | Cluster6 |
| *OsTN9g001610* | *bHLH130* | Cluster6 |
| *OsTN9g001682* | *bHLH089* | Cluster6 |

**Table S4.**
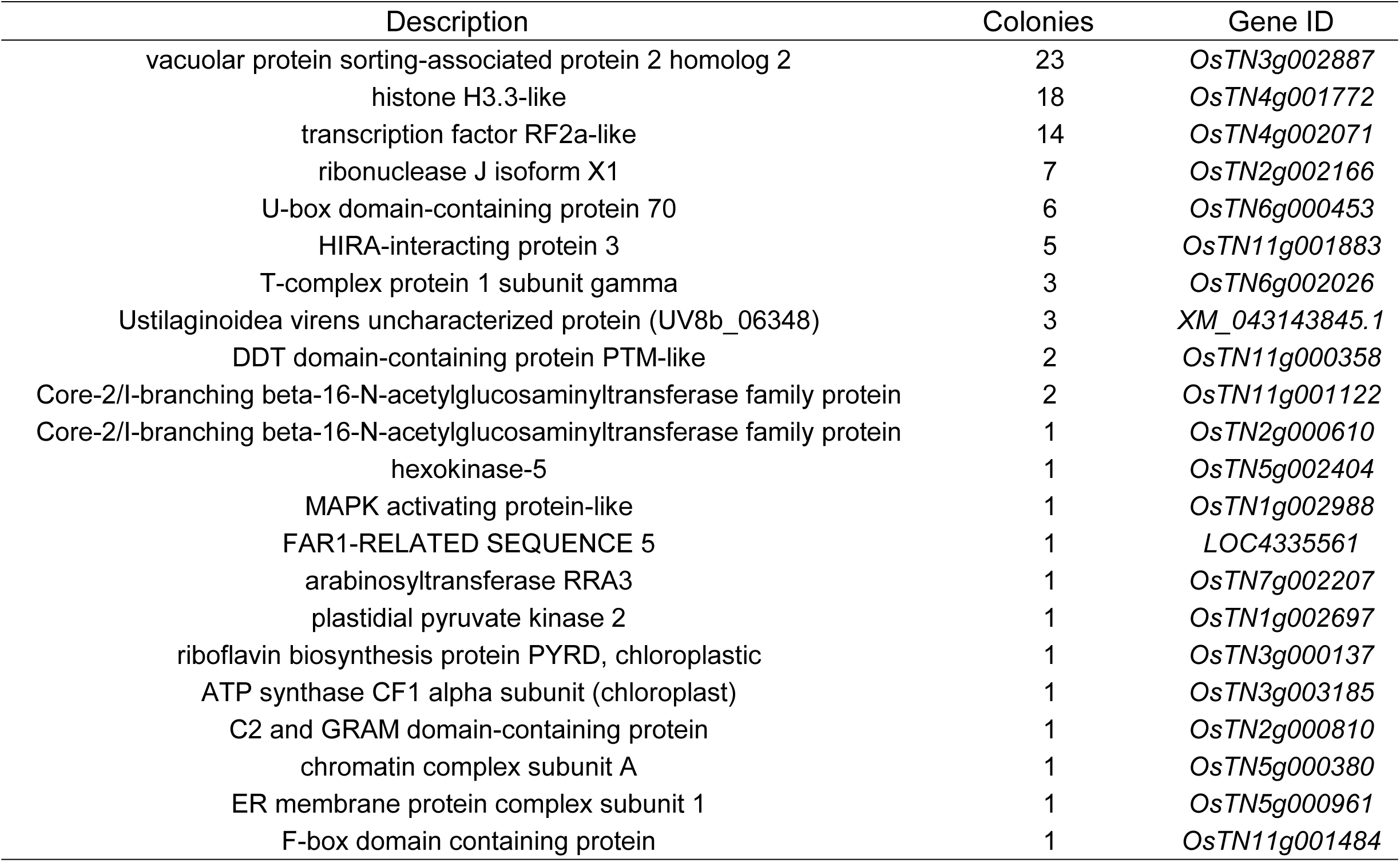
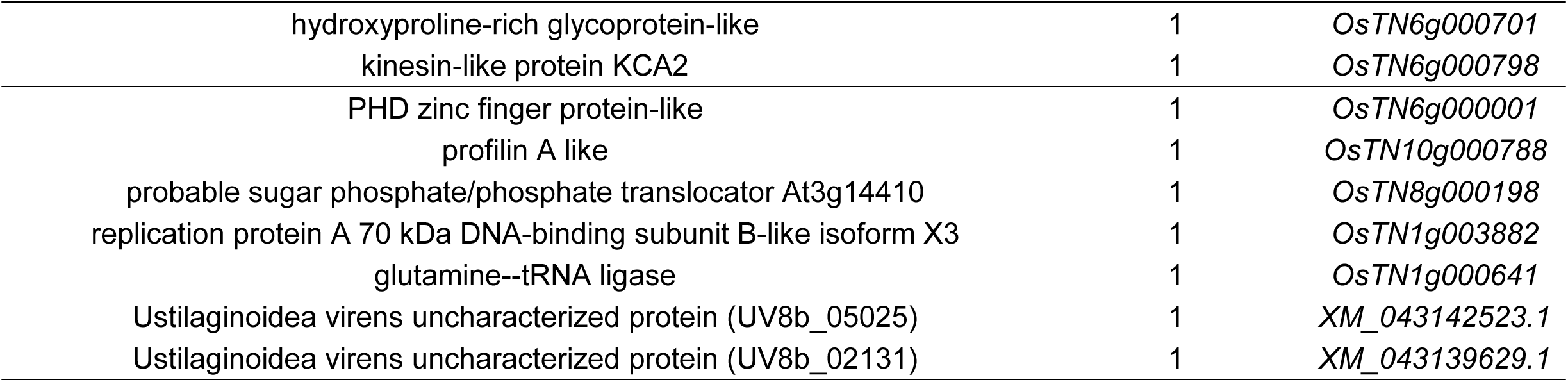
Statistics on the number of monoclonal colonies for each potential *BSCL1*-interacting protein. The raw data of screened out monoclonal colonies via yeast three-hybrid are listed in Dataset S6.

**Table S5.**
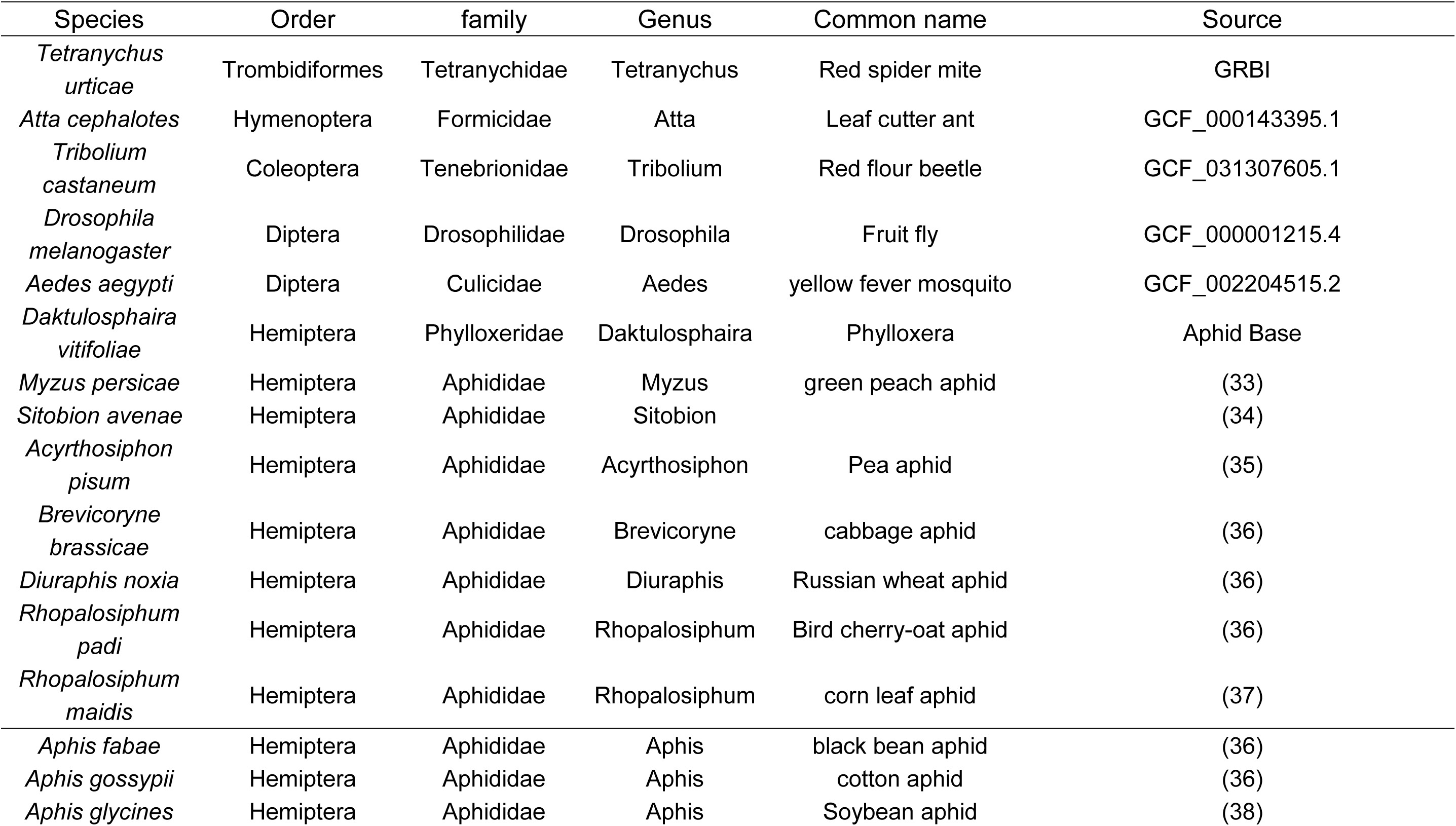
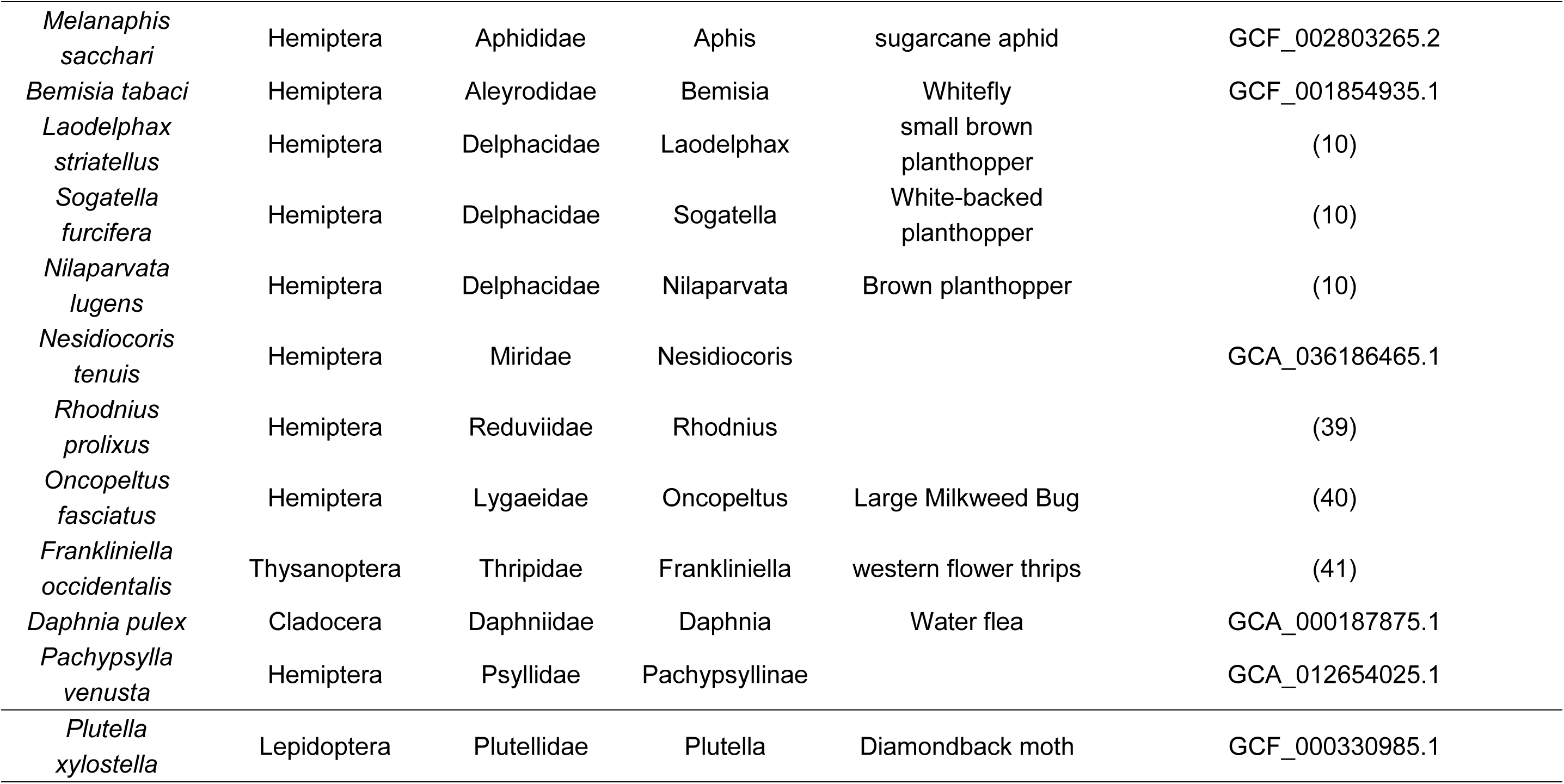
Genome data source of 28 species used in Evolinc Ⅱ analysis.

**Table S6.**
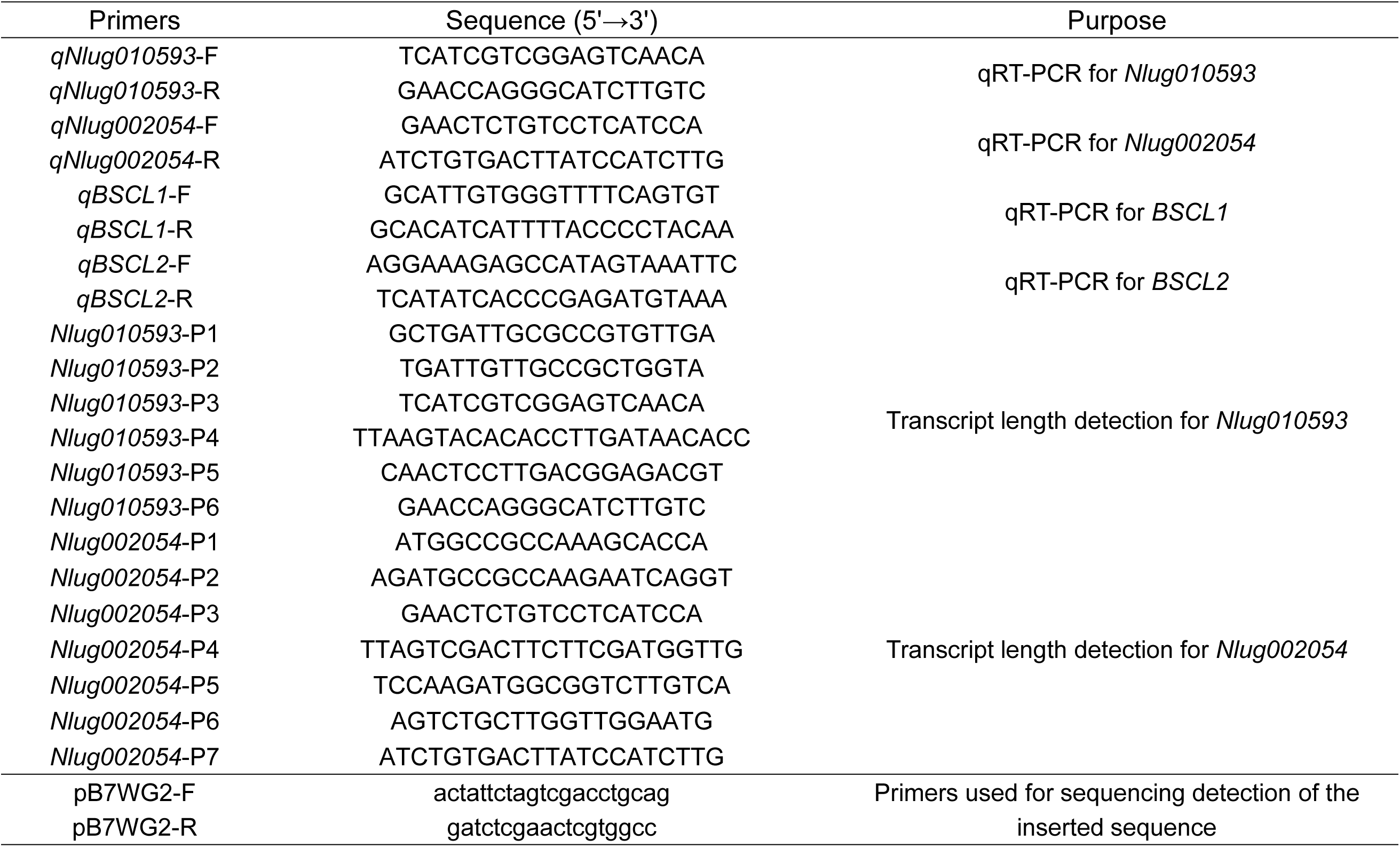
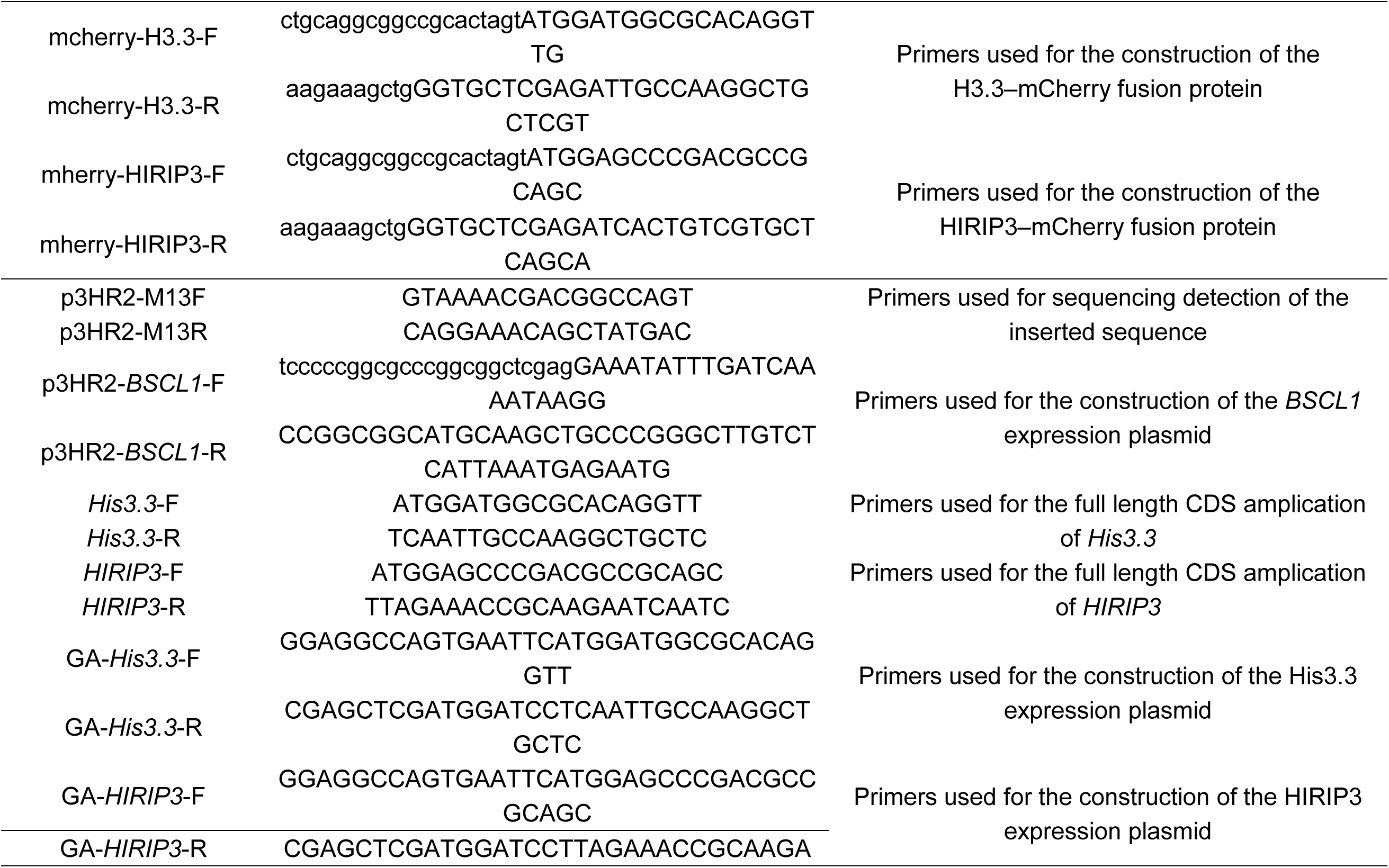
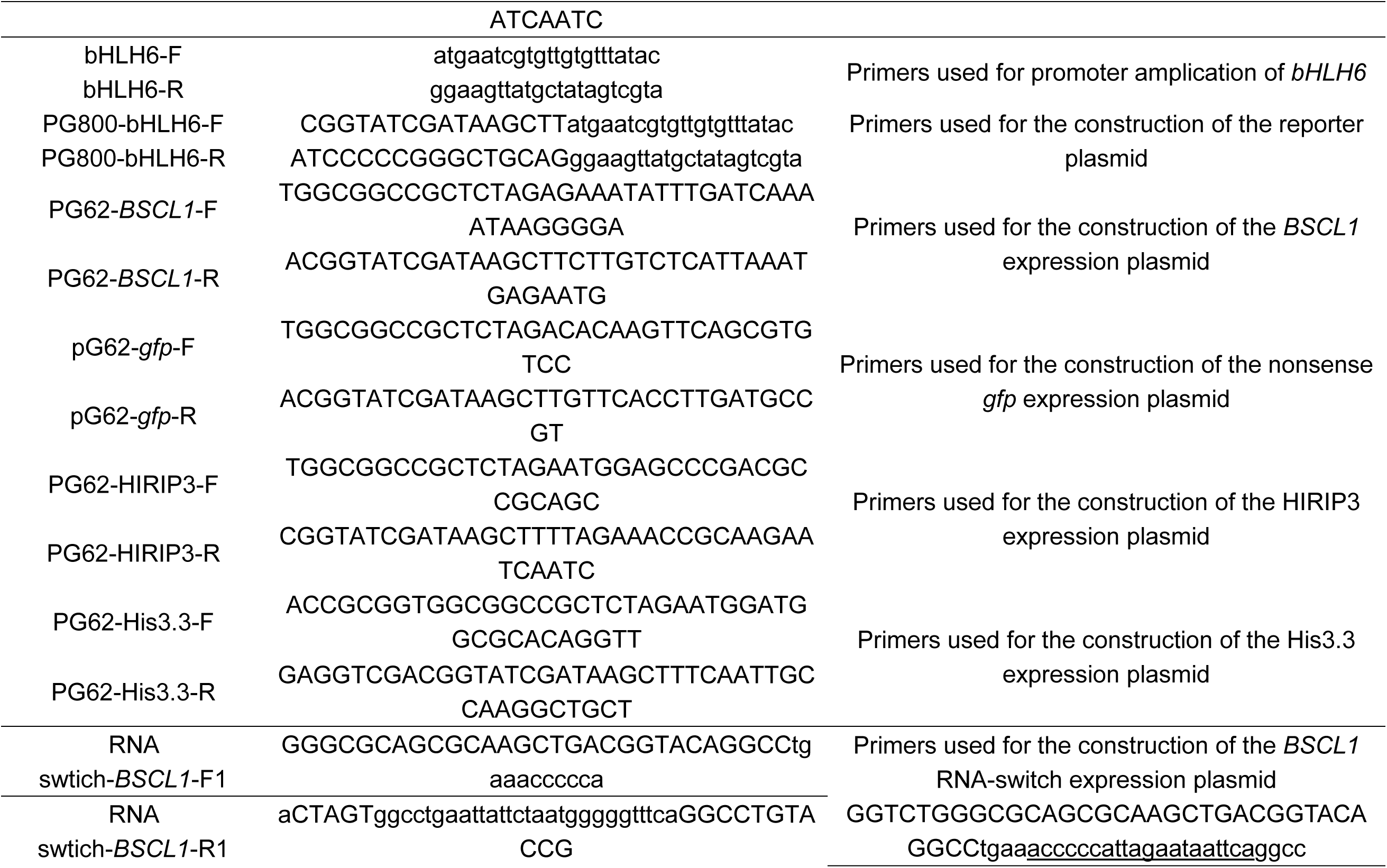
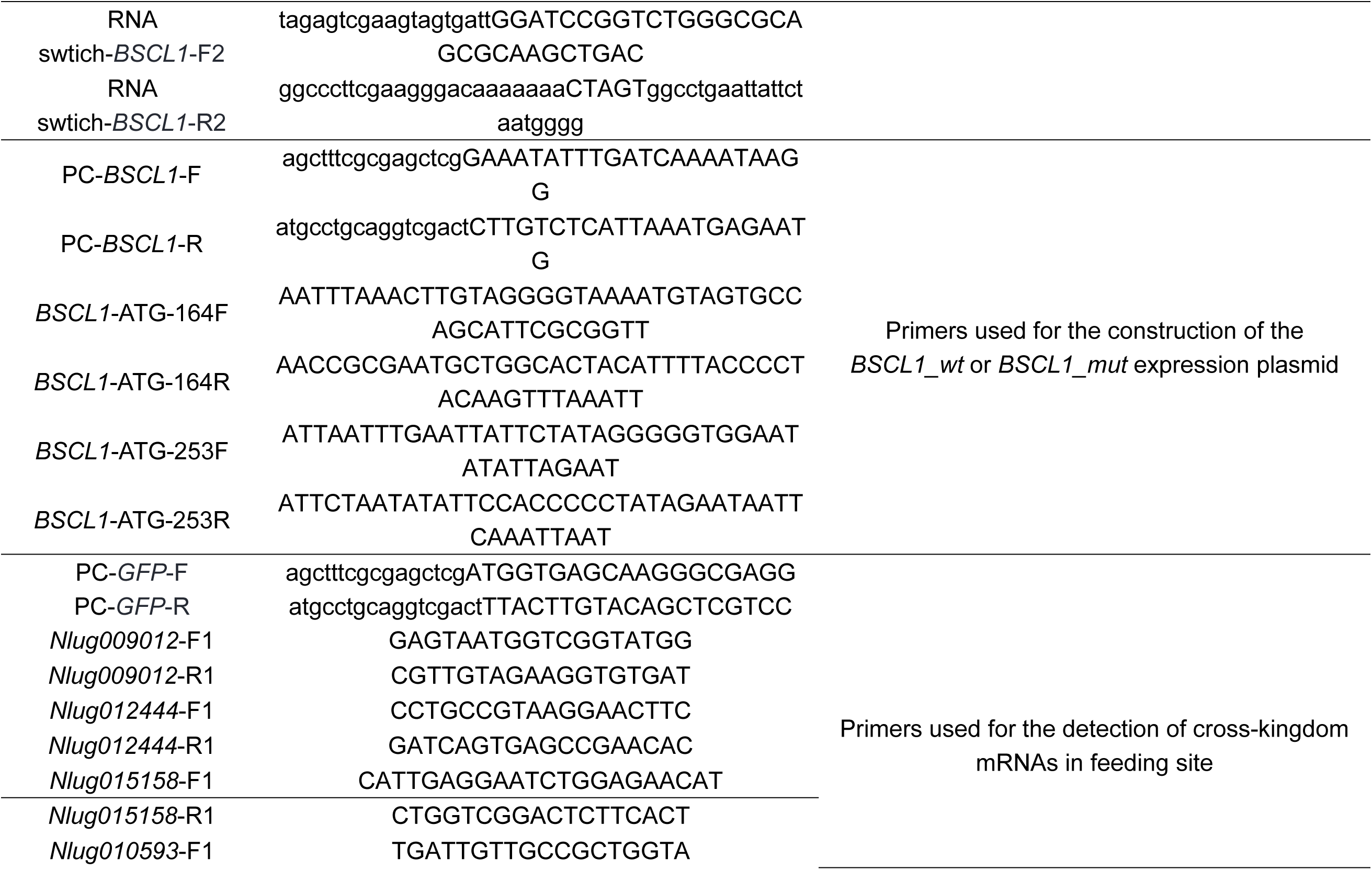
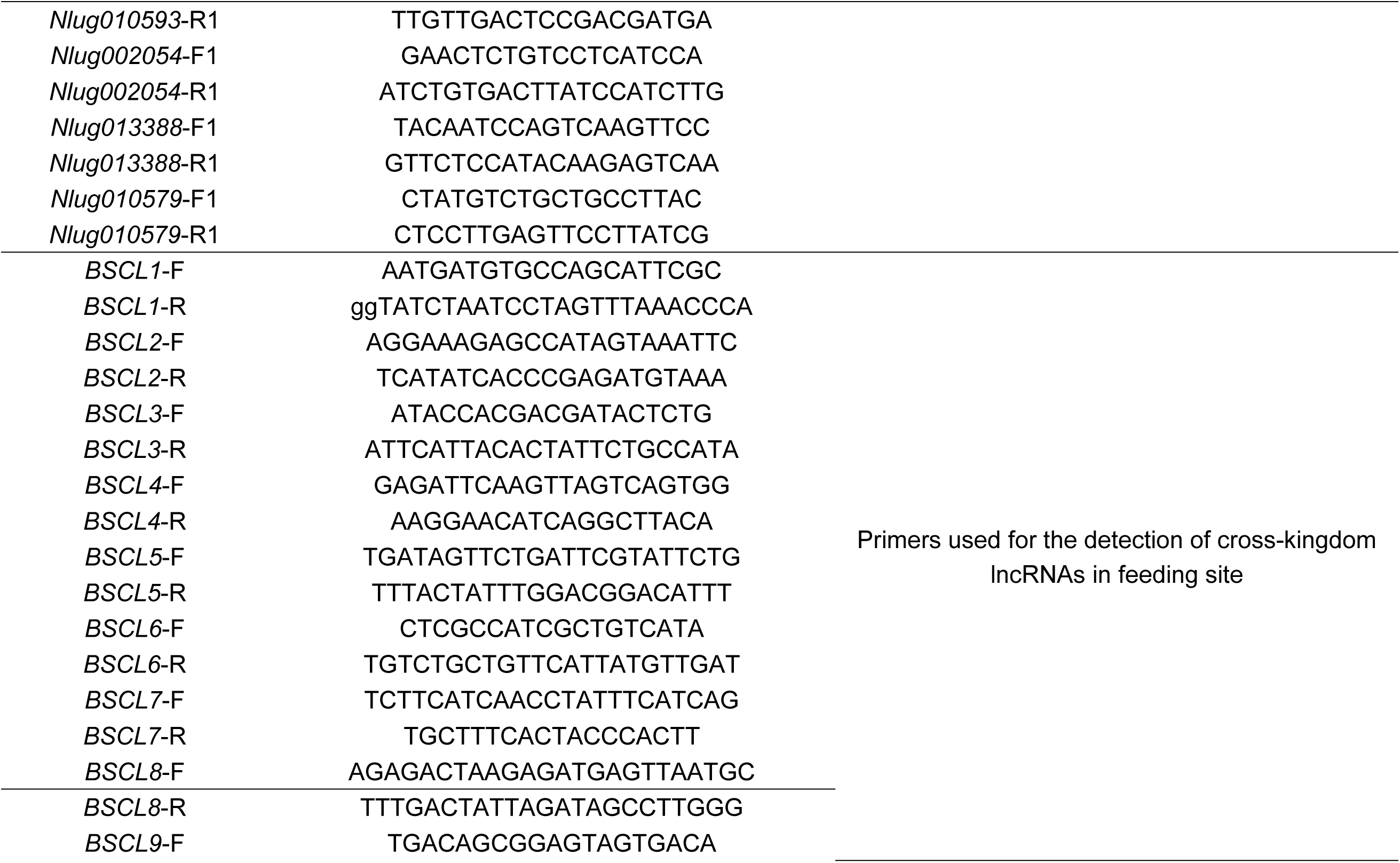
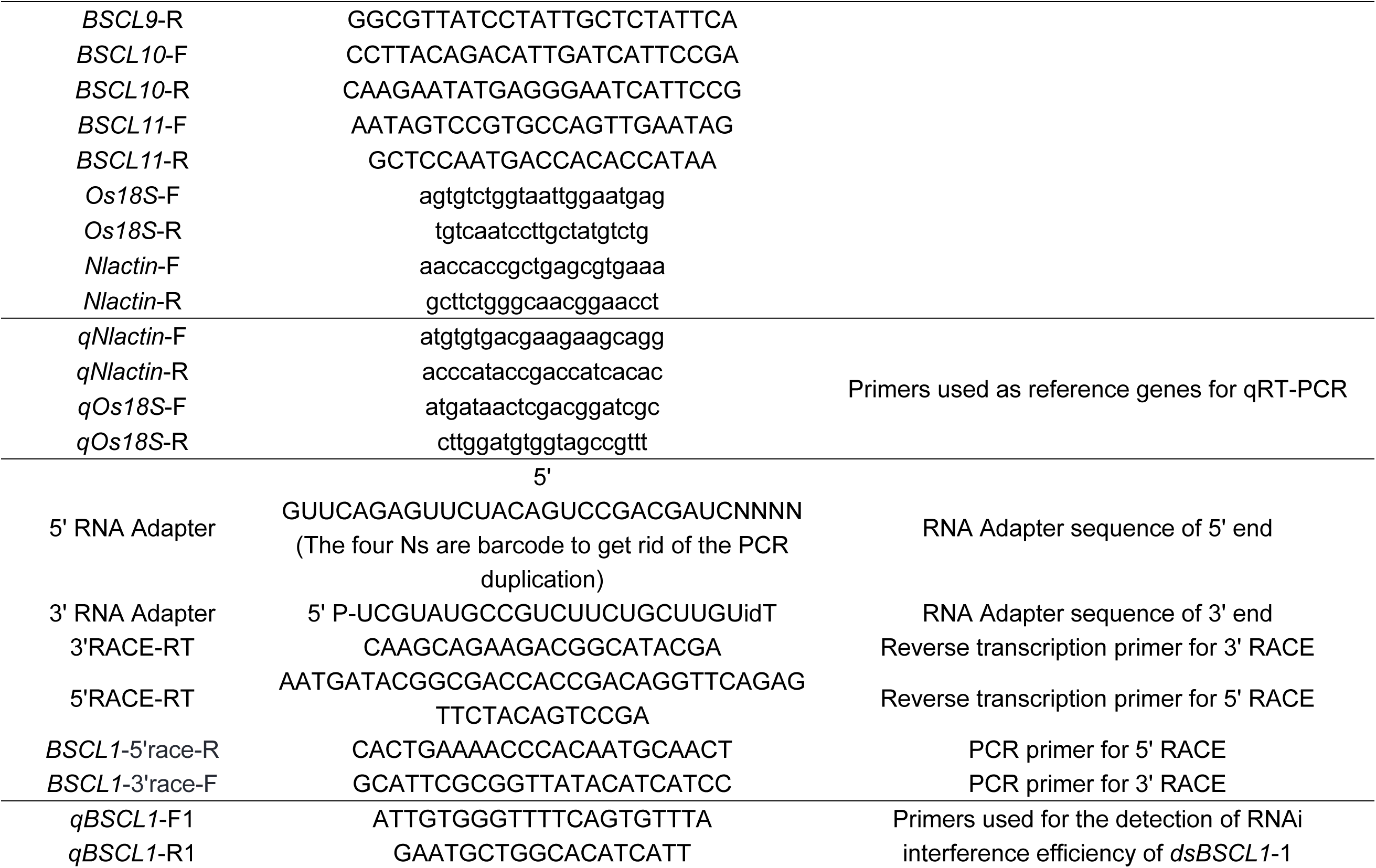
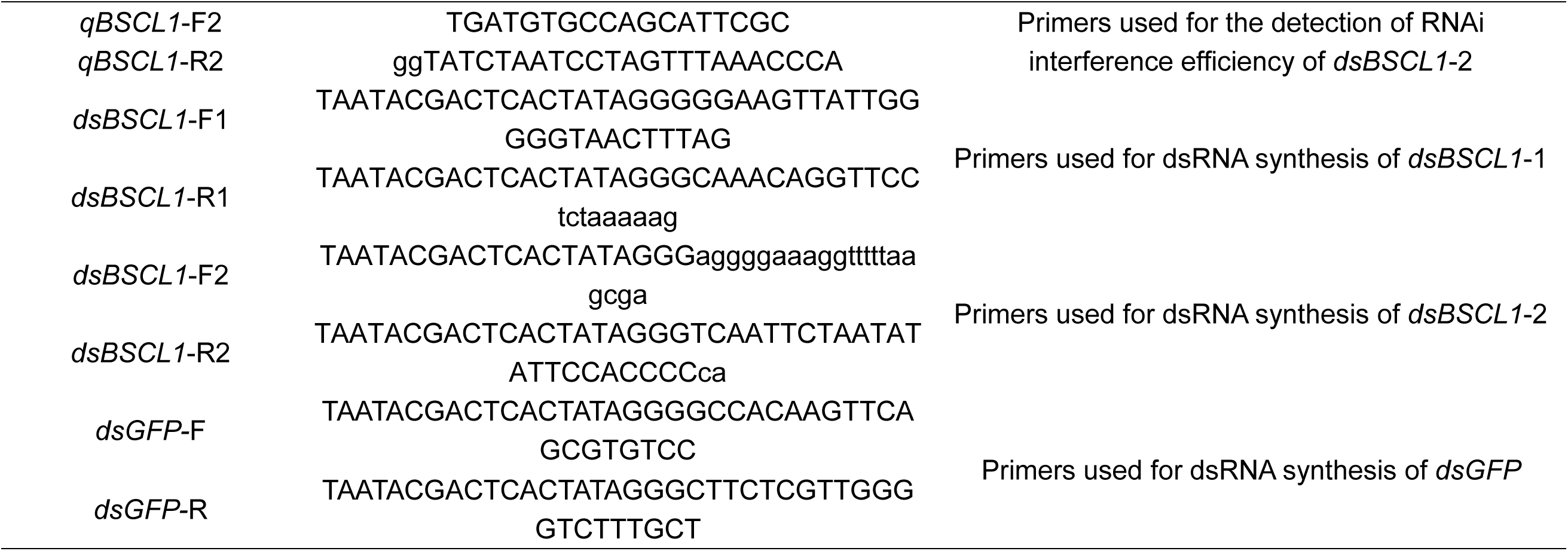
Primers used in this study.

## Legends for Datasets S1 to S6

**Dataset S1: List of differentially expressed genes of BPH feeding on resistant rice RHT and susceptible rice TN1.**

**Dataset S2: Statistics of analyses of RNA-seq data retrieved from BPH-exposed (feeding sites) and non-exposed (control) rice sheaths. RNA-seq reads derived from 2 to 3 biological replicates per treatment were mapped to TN1 rice and BPH merged genome.**

**Dataset S3: Reads count of identified cross-kingdom RNAs in sample BPH-1-FS from replicate 1.**

**Dataset S4: List of differentially expressed genes in different rice lines with or without BPH feeding.**

Table 1. Differentially expressed gene sets in TN1 rice with and without BPH feeding.

Table 2. Differentially expressed gene sets in over-expression GFP TN1 rice with and without BPH feeding.

Table 3. Differentially expressed gene sets in over-expression *BSCL1_wt* TN1 rice with and without BPH feeding.

Table 4. Differentially expressed gene sets between TN1 and over-expression *BSCL1_wt* TN1 rice.

Table 5. Differentially expressed gene sets between TN1 and over-expression *BSCL1_wt* TN1 rice with BPH feeding.

Table 6. Differentially expressed gene sets between over-expression GFP and over-expression *BSCL1_wt* TN1 rice.

Table 7. Differentially expressed gene sets between over-expression GFP and over-expression *BSCL1_wt* TN1 rice with BPH feeding.

**Dataset S5: List of differentially expressed genes and their expression patterns in different clusters.**

**Dataset S6: List of potential *BSCL1*-interacting proteins in rice screened out vie yeast three-hybrid assays**

